# Deep embedding and alignment of protein sequences

**DOI:** 10.1101/2021.11.15.468653

**Authors:** Felipe Llinares-López, Quentin Berthet, Mathieu Blondel, Olivier Teboul, Jean-Philippe Vert

## Abstract

Protein sequence alignment is a key component of most bioinformatics pipelines to study the structures and functions of proteins. Aligning highly divergent sequences remains, however, a difficult task that current algorithms often fail to perform accurately, leaving many proteins or open reading frames poorly annotated. Here, we leverage recent advances in deep learning for language modelling and differentiable programming to propose DEDAL, a flexible model to align protein sequences and detect homologs. DEDAL is a machine learning-based model that learns to align sequences by observing large datasets of raw protein sequences and of correct alignments. Once trained, we show that DEDAL improves by up to two- or three-fold the alignment correctness over existing methods on remote homologs, and better discriminates remote homologs from evolutionarily unrelated sequences, paving the way to improvements on many downstream tasks relying on sequence alignment in structural and functional genomics.

## Introduction

Sequence alignment is a key component of bioinformatics pipelines to study the structures and functions of proteins, and to annotate open reading frames (ORF) in newly sequenced genomes and metagenomes [1]. Indeed, aligning two protein sequences allows identifying homologous sequences with known structure or function, and regions of similarity that may be the result of evolutionary, structural or functional relationships. Jointly aligning multiple sequences further gives information about evolutionary constraints and patterns of co-evolution along the sequence, which is for example useful for 3D structure inference [2–4]. It is remarkable that algorithms for pairwise sequence alignment have basically remained unchanged since their invention in the 1980’s and 1990’s, in particular the Smith-Waterman (SW) algorithm to find the best local alignment between two sequences in quadratic time with dynamic programming [5] and faster heuristics like BLAST [6] or FASTA [7] which are also the first steps of more advanced methods based on multiple sequence alignment such as PSI-BLAST [8]. However, in spite of their immense popularity and importance for downstream applications, these algorithms often produce erroneous alignments or fail to detect homology, particularly for sequences with low similarity [9]. This leaves a sizeable fraction of predicted ORF in genomics and metagenomics projects without annotation [10], while alignment errors can in turn result in wrong structural or functional annotations. Improving pairwise sequence alignment algorithms, particularly for divergent sequences, could directly benefit a number of downstream tasks.

The SW algorithm formulates the problem of finding the correct alignment between two sequences as a search for the best-scoring path in a graph. Each candidate path represents a possible alignment and the score of a path depends on user-defined parameters. These are the costs of starting or extending a gap in the alignment, and the substitution score of aligning each position of the first sequence to each position of the second one, which usually depends on the amino acids at both positions. This formulation, which leads to efficient algorithms to find or approximate the best-scoring alignment, has at least two weaknesses that may lead to possible errors in the alignment found. First, it lacks flexibility in the formulation of the score of a path, by enforcing for example that the cost of opening or extending a gap is the same wherever in the sequences, or by enforcing that the substitution score of aligning two positions only depends on the amino acids at those positions. Allowing more flexible parameterizations may give more opportunities to the algorithm to optimize a relevant score; for example, [11] show the benefits of making the substitution score between two positions in protein sequences depend not only on the amino acids at those positions, but also on predicted structural properties, at the cost of increasing the number of parameters of the model. Second, the best alignment returned by the algorithm strongly depends on the choice of the parameters [12–18], and even for simple models where the substitution score only depends on the amino acids to be aligned, it is well known that there is no universally “good” substitution scores that lead to good alignments for all protein pairs [19, 20]. This raises the question of how to optimally choose adequate scoring parameters for a given pair of proteins.

In this work, we propose DEDAL (*Deep Embedding and Differentiable ALignment*), an algorithm for pairwise sequence alignments that addresses both issues. DEDAL builds on top of the standard SW algorithm to efficiently find an optimal alignment between two sequences, but provides a flexible parameterization of the scoring function used by the SW algorithm that adapts to each sequence pair and each position in each sequence. The parameterization is automatically learned during a training phase from a set of sequence pairs with known alignments, and a large set of raw protein sequences. It relies both on recent advances in deep learning language models which embed discrete sequences in a continuous space and are automatically trained on a large corpus of raw sequences, and also a parameterization of the SW algorithm (gap and substitution parameters) as a function of the continuous embedding. To train DEDAL, we propose a smoothed variant of the SW algorithm, to make the alignment solution a continuous and differentiable function of the scoring parameters. Given a set of sequence pairs with known correct alignment, we then tune automatically the various parameters of the model by end-to-end gradient-based optimization to minimize the alignment errors. Once trained, DEDAL produces gap and substitution scoring matrices computed specifically for each new pair of sequences. In addition, the gap and substitution scores are contextual: for each pair of positions, they depend on the full sequences to be aligned. The optimal alignment is then computed with a standard SW algorithm using those parameters. We show that DEDAL can be trained efficiently on modern hardware with accelerators. Once trained, we demonstrate that DEDAL improves by up to two- or three-fold the quality of the alignment predicted for remote homologs compared to standard SW, and produces an alignment score that more accurately detects remote homology.

### Related work

The fact that the solution of the SW algorithm depends on the scoring parameters has been studied in detail in the context of the so-called parametric sequence alignment problem, which aims to describe the set of possible solutions as a function of the parameters used [12–18]. This approach, however, is only feasible for simple models with up to 2 or 3 parameters that vary. Conversely, the idea to search for parameters that reproduce a set of given, correct alignments, was tackled by [11, 21–23] using various optimization techniques, but these works focus on simple parameterizations where a fixed-size substitution matrix and one or two gap parameters are optimized. It is proposed in [24] to exploit known alignments to help train deep embedding models with a differentiable loss, however this model does not produce a sequence alignment once trained. The closest work to ours is the DeepBlast model of [25], who also propose to learn a deep language model for proteins with a differentiable alignment module, however the model has a simpler model for scores (linear instead of affine), and only produces global alignments. More recently, concurrently and independently from this work, [26] presented a differentiable version of SW for multiple sequence alignment, albeit with a simple convolution layer instead of a full language model to embed the sequences, and simpler position-independent gap penalty. Moreover, they focus on contact and structure prediction tasks, as opposed to directly investigating alignment and homology detection performance as done here.

## Results

### Pairwise local alignment of protein sequences with DEDAL

We introduce DEDAL, a trainable algorithm for accurate pairwise local alignment of protein sequences (Figure 1). DEDAL aligns sequences by computing substitution scores and gap penalties that are specific to the sequences being aligned (Figure 1, top). To this end, DEDAL depends on parameters which are automatically tuned before inference during a training phase (Figure 1, bottom).

**Figure 1:**
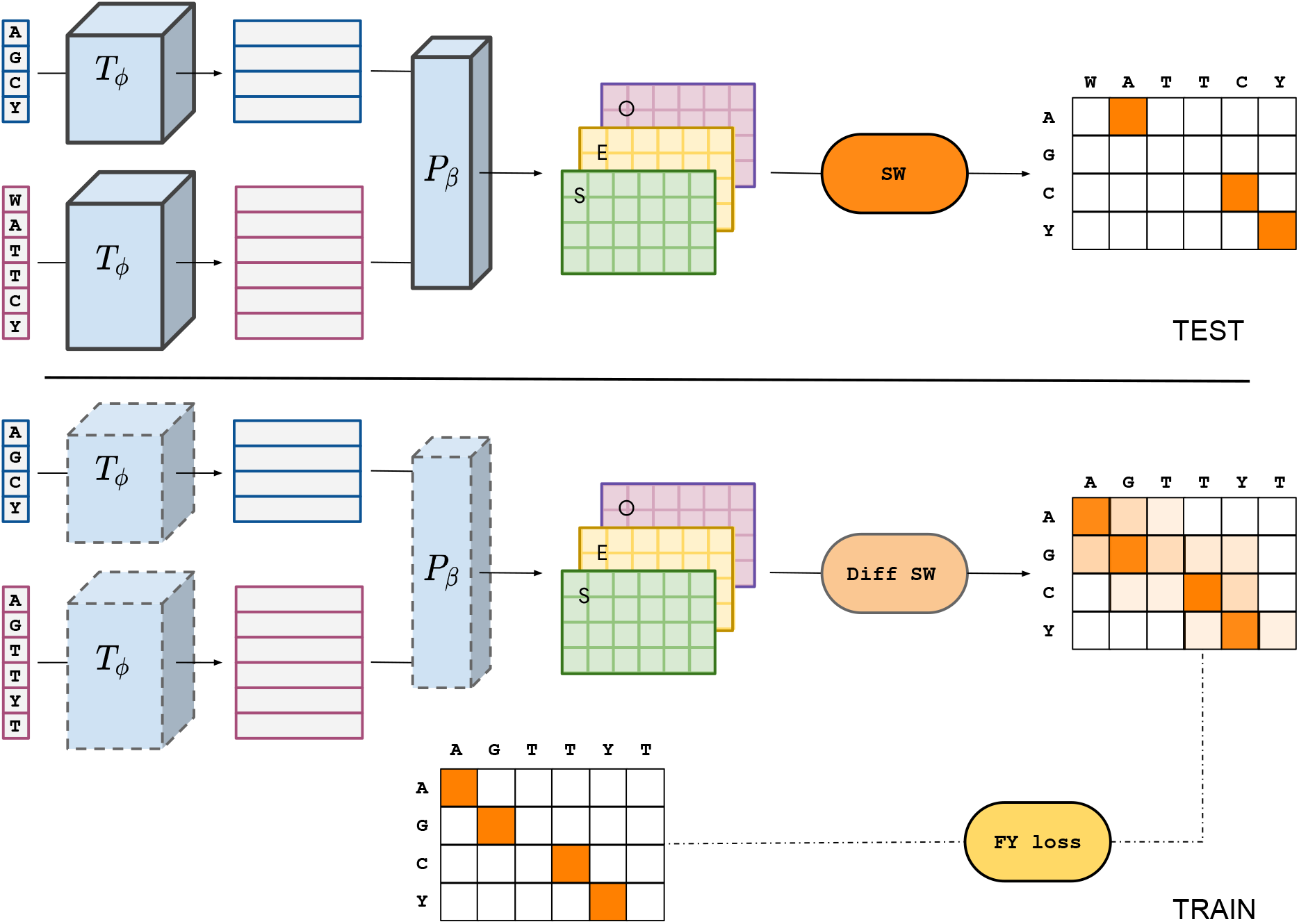
Overview of DEDAL. At test time (top), DEDAL aligns two sequences *x* and *y* by running the SW local alignment algorithm with position-specific parameters (gap open *O*, gap extend *E*, substitution scores *S*) that depend on the input sequences through a transformer encoding *T_ϕ_* of each sequence followed by a parameterizer *P_β_* that transforms the continuous representations into matrices of parameters needed by SW. Both *T_ϕ_* and *P_β_* depend on parameters *ϕ* and *β* which are learned at train time from a large corpus of raw sequences to learn the language model *T_ϕ_*, combined with pairs of sequences with known alignment to learn jointly *T_ϕ_* and *P_β_* by end-to-end optimization with a differentiable variant of SW and a specific alignment loss function (bottom).

For sake of clarity, let us first describe how DEDAL works once it is trained and ready to be used to align sequences. In order to align two sequences *x* and *y* and score the resulting alignment, DEDAL simply uses the standard SW algorithm for pairwise local alignments, but with gap open, gap extend and substitution score matrices which are computed specifically from *x* and *y*. To do so, a deep learning-based transformer encoder network *T_ϕ_* with parameters *ϕ* [27] is first used to independently obtain a continuous representation of each of these sequences. In this representation, each residue of each sequence is mapped to a vector in a vector space of a fixed dimension (we use *d* = 768 in our experiments). Crucially, these embeddings are *contextual*, that is, the embedding of each residue encodes information not only about the amino acid present at that position but also about all other residues in the sequence, as well as their relative arrangement. This allows DEDAL to be highly flexible in the way sequences are represented, opting for a data-driven approach towards incorporating contextual information over hard-coded rules. Next, DEDAL computes a substitution score as well as gap open and gap extend penalties for each pair of residues from the sequences to be aligned, computed from their respective vector representations by a parameterizer function *P_β_*, which depends on parameters *β*. Finally, the standard SW algorithm is used to compute an optimal alignment and score it, using the substitution scores, gap open and gap extend penalties computed at the previous step. In other words, DEDAL relies on the SW algorithm to align sequences and score the alignment, but provides a very flexible framework to parameterize the SW algorithm; in particular, the substitution score, gap open and gap extend penalties are specific to each pair of positions in the two input sequences, and depend on the full sequences through the contextual embedding of the transformer encoder and the parameterizer.

DEDAL thus depends on parameters *ϕ* for the transformer encoder *T_ϕ_*, and *β* for the parameterizer *P_β_*. These parameters are tuned automatically during a training phase, where we provide DEDAL with a large set of ~ 30 million non-redundant protein sequences from UniRef50 [28] to train the transformer encoder, as well as a set of pairs of homologous sequences with curated correct alignment extracted from the set of ~ 1.2 million sequences organized in ~ 19 thousand families in the Pfam-A seed database [29] to jointly train the transformer encoder and the parameterizer. More precisely, using these datasets, DEDAL tunes its parameters by end-to-end gradient-based optimization to minimize a loss function combining three tasks: 1) a so-called *masked language modelling* task on the UniRef50 dataset, which is a classical way to tune the parameters of a transformer encoder [30, 31]; 2) an *homology detection* task, where we train DEDAL so that it can discriminate homologous from non-homologous pairs in Pfam-A seed from the alignment score; 3) a *learning to align* task, where we train DEDAL to produce the correct alignment on homologous pairs from Pfam-A seed with known correct alignment. In order to solve the second and third tasks by gradient-based optimization, we implement a novel differentiable variant of the SW algorithms that allows to backpropagate not only the alignment score but also the error between the predicted and true alignments to the parameters *ϕ* and *β*. Since Pfam sequences represent protein domains, which therefore tend to be aligned from start to end within a given family, we perform data augmentation, *extending* these sequences by including flanks upstream and downstream of the domain so that DEDAL is trained to produce a correct *local* alignment. We refer the reader to the Methods section for more details about the model and the data. DEDAL is implemented in Python, using the TensorFlow [32] library for deep learning. It takes about one week using 32 Tensor Processing Unit (TPU) v3 cores to train DEDAL.

### DEDAL accurately aligns homologous sequences

We first assess the ability of DEDAL to accurately align homologous sequences. Since DEDAL is trained on a set of known correct alignments, we must evaluate its performance on sequences not seen at train time. We therefore split the Pfam sequences into two non-overlapping sets. The first one is used to train the model and to choose hyperparameters, and the second one to assess its performance. We consider two ways to make the split: 1) split sequences uniformly at random in Pfam, so that sequences in the test set come from families seen at train time (we call this setting the *in-distribution* setting), and 2) split clans between training and test sets, so that sequences in the test set come from clans that are not seen at train time (we call this setting the *out-of-distribution* setting). Obviously the out-of-distribution setting is more challenging, since it requires the model to generalize across clans. It mimics the situation where we want to align sequences from unknown domains. The in-distribution setting, on the other hand, mimics the common situation where we want to align sequences from known Pfam domains. In both cases we keep the UniRef50 set of sequences used for the masked language model task, since we want to simulate the situation where a user wants to align a sequence from the “protein universe” described by UniRef50 and known at training time, whether or not it is similar to sequences annotated in Pfam; in addition, we provide in Supplementary Section S2.1 more results where we only train the language model task on a subset of UniRef50 with no match to any of the Pfam families in the out-of-distribution test set to assess DEDAL’s performance on sequences that are not only in Pfam clans not seen at training time, but are also significantly different from any sequence in the “protein universe” seen at training time for the language model task. We measure the quality of a predicted alignment by the *F*_1_ score of the prediction on the test set, and stratify the performance by percent identity (PID) of the sequences in the correct alignment, since it is well known that the difficulty of aligning sequences increases when PID decreases. As a baseline, we compare DEDAL to the SW algorithm ran with a fixed PFASUM70 substitution matrix and an affine gap penalty model with gap open=15 and gap extend=1.5. We chose this configuration as the best among more than 1,400 combinations of substitution matrices in the BLOSUM [33], VTML [34, 35] and PFASUM [20] families, and gap open and extend parameters (see Methods).

Figures 2a and 2b summarize the alignment performance of DEDAL and of the baseline, respectively in the in-distribution and out-of-distribution settings. A breakdown of the *F*_1_ score into precision and recall is shown in Supp. Figures S8 and S9, respectively. We see that DEDAL significantly outperforms the baseline both in-distribution and out-of-distribution. The difference is particularly strong in-distribution (with an average relative improvement of 18% over baselines, from *F*_1_ = 0.744 to *F*_1_ = 0.877), which confirms the benefit of training DEDAL on Pfam families in order to then align new protein sequences within these families. Strikingly, the gap between DEDAL and baselines widens as the PID decreases, with a relative improvement of up to 286% in the hardest setting (PID ≤ 0.1), showing that DEDAL is able to provide good quality alignments (*F*_1_ = 0.587) even between very remote homologs, while the baseline can not (*F*_1_ = 0.152). Interestingly, a similar pattern is visible on the out-of-distribution setting, where DEDAL outperforms the baselines by 4% on average (from *F*_1_ = 0.782 to *F*_1_ = 0.816), and by 119% on the remote homologs with PID < 0.1 (from *F*_1_ = 0.150 to *F*_1_ = 0.329). This suggests that DEDAL learns a model of protein sequence similarity that extends beyond the biological subspace it sees at train time, and that DEDAL is particularly useful is difficult situations of remote homology, where standard methods perform poorly.

**Figure 2:**
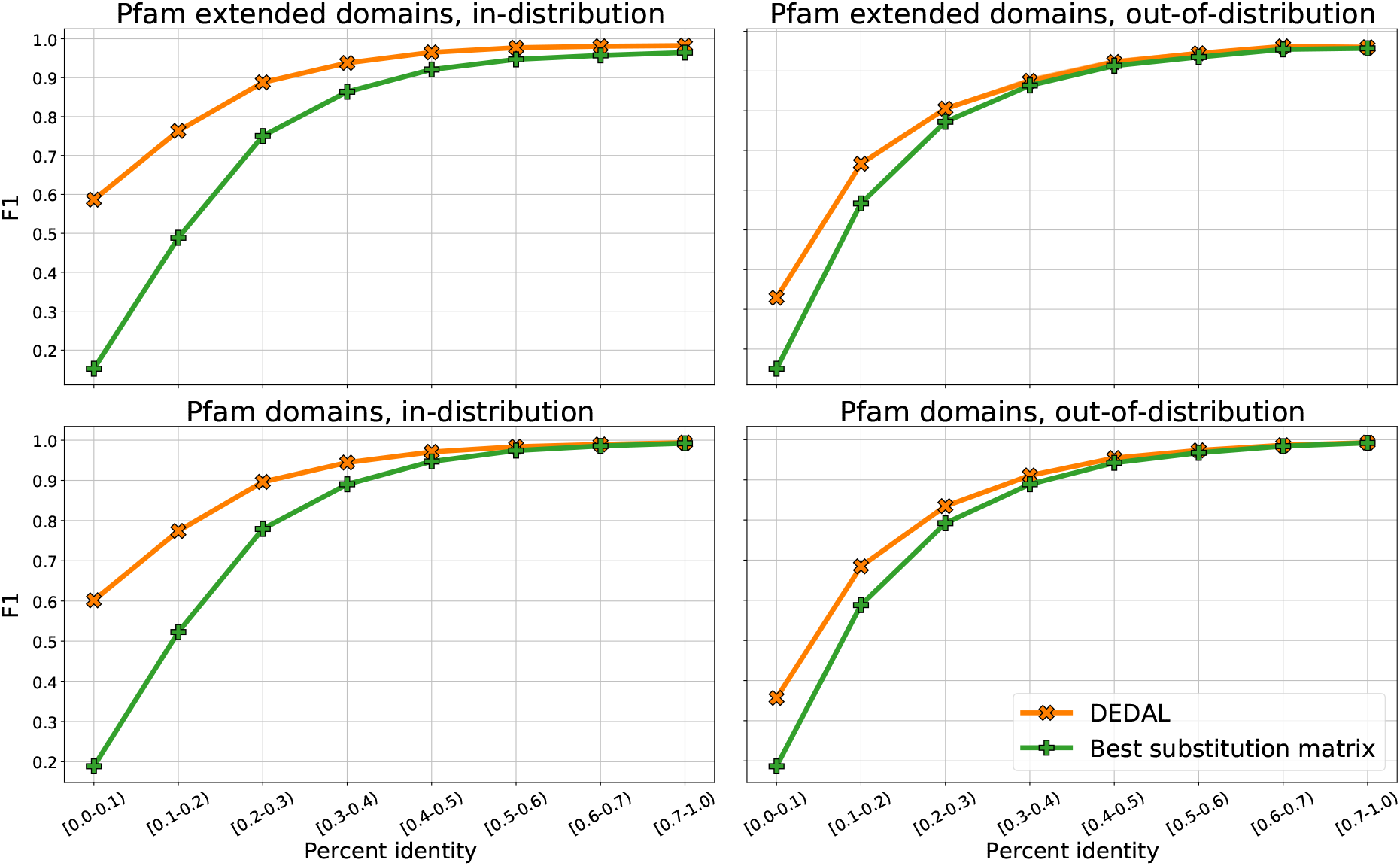
Alignment *F*_1_ score of DEDAL and the best-performing substitution matrix baseline, in the in- and out-of-distribution settings (respectively, left and right columns), and for Pfam extended or raw domains (respectively, top and bottom rows).

We then compare DEDAL and the baseline on their ability to align Pfam domains, as opposed to the extended domains that we use to train DEDAL. While the domain sequences are the same in both cases, the alignments are very different since two domain sequences in a Pfam family can generally be aligned from the beginning to the end, thus making the correct alignment closer to a global alignment than a local one. We show the results in this setting in Figures 2c and 2d. Compared to the previous setting, we see that the performances of all methods tend to be better, which can be explained by the fact that the number of wrong candidate alignments is reduced when we focus on the domain sequence that must be aligned. Besides the overall improvement of all methods, DEDAL still significantly outperforms the baseline, particularly in-distribution and particularly in the low PID regime (with, e.g., a relative *F*_1_ increase of 218% for in-distribution domains with PID ≤ 0.1, from *F*_1_ = 0.189 to *F*_1_ = 0.602).

In order to clarify why DEDAL behaves so differently from a standard SW algorithm with a fixed substitution matrix and gap parameters, we now focus on a particular pairwise alignment between two domain sequences of Pfam-A seed (accession PF00048): a C-C motif chemokine from western lowland gorilla (UniProt A0A2I2ZUR5) and a C-X-C motif chemokine ligand 14 from northern mallard (UniProt U3IBZ2). Figure 3 summarizes the alignments produced by DEDAL and the baseline (using a PFASUM substitution matrix), and the parameters of both models. Note that this alignment is not part of DEDAL’s training set. As seen in Figure 3.a, the sequences have only 8 conserved residues in the Pfam-A seed curated alignment, meaning that this is a difficult case (PID=0.12). Indeed, using SW with a PFASUM substitution matrix gives an alignment where no residue is correctly aligned (*F*_1_ = 0), however we see that using DEDAL gives an almost perfect alignment where only 4 residues are wrongly aligned, and in all cases with an error of only one residue in distance (*F*_1_ = 0.94). To better understand the difference between PFASUM and DEDAL, we plot in Figure 3.b and 3.c, respectively, the SW parameters for PFASUM (substitution scores) and for DEDAL (substitution scores, gap open and gap extend penalties). We see that the substitution score matrix produced by DEDAL is much smoother spatially than the one of PFASUM, highlighting the benefit of context dependent embeddings of residues. We also see that DEDAL clearly picks the importance of aligning cysteine (C) residues, which are very conserved since they form disulfide bonds which stabilize the 3D structure of the domain (there are four cysteines in each sequences, respectively at positions 3, 4, 27, 43 in gorilla and 3, 5, 29, 49 in mallard, corresponding to the 16 positions in DEDAL’s substitution matrix with highest score). PFASUM, on the other hand, has high scores not only to match the cysteines, but also to match the two tryptophans (W) at respective positions 21 and 49 in the gorilla sequence, and 38 and 63 in the mallard sequence. While tryptophans are usually very conserved, and have therefore a large substitution score in standard substitution matrices such as PFASUM, it is interesting that in this particular case they should not be aligned according to the Pfam-A seed curated alignment, and DEDAL correctly predicts it by producing a small substitution score. Finally, we note that DEDAL’s gap open and extend penalties are far from uniform in the matrix, with in particular strong penalties to open and extend gaps near the cysteines, suggesting that DEDAL has learned that mutations are more likely than insertions or deletions near a cysteine involved in a disulfide bond.

**Figure 3:**
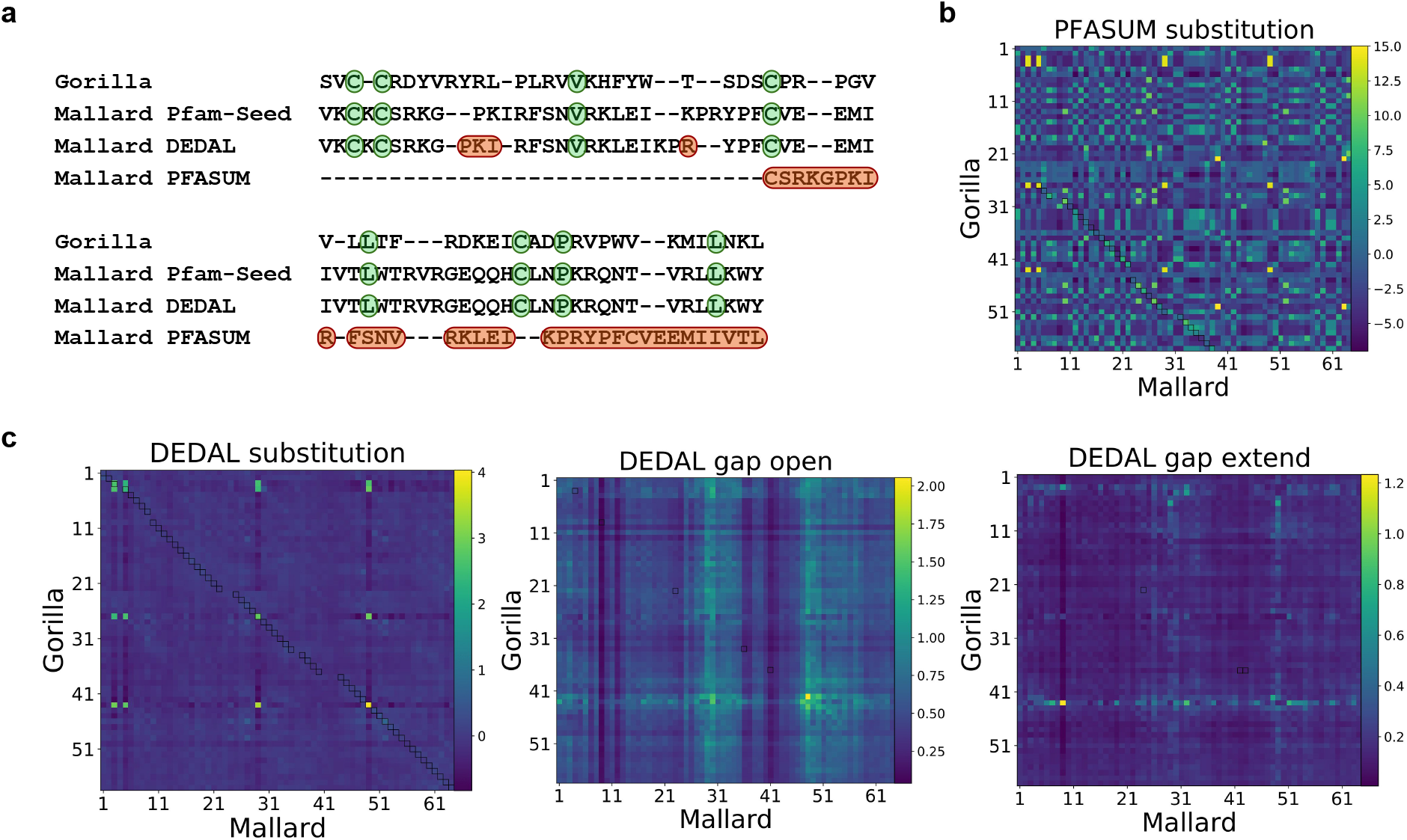
Example of pairwise alignment of two protein domain sequences from Pfam-A seed, a C-C motif chemokine from western lowland gorilla (UniProt A0A2I2ZUR5) and a C-X-C motif chemokine ligand 14 from northern mallard (UniProt U3IBZ2). **a.** Alignment respectively from the curated Pfam-A seed database (second row), predicted by DEDAL (third row), and predicted with a PFASUM70 substitution matrix (fourth row). We show all residues in both sequences for the Pfam-A seed and DEDAL alignments, but not the unaligned residues upstream and downstream of the alignment in the mallard sequence for PFASUM. Residues highlighted in green correspond to correctly aligned conserved residues, while those in red correspond to discrepancies between a predicted alignment and the Pfam-A seed one. **b.** Substitution scores between all pairs of residues from the PFASUM substitution matrix. **c.** SW parameters predicted by DEDAL.

### DEDAL accurately detects remote homologs

Next, we seek to determine if DEDAL’s ability to align homologous sequences accurately also implies that the alignment score it computes is effective to detect homology. To this end, we follow the same experimental setup as when probing alignment performance, namely considering pairs of Pfam extended domains or pairs of Pfam domains, on the one hand, and considering pairs of candidate homologous sequences from families seen at train time (in-distribution setting) or from clans not seen at train time (out-of-distribution setting), on the other hand. For each of these settings, we measure the area under the ROC curve (AUROC) when predicting whether a pair of sequences belongs to the same Pfam clan or not based on the SW score, correcting the alignment score for sequence length as usually done when computing alignment E-values to assess homology (see Methods). To rank true homologs according to the “difficulty” of detecting their relatedness, those originating from the same Pfam family are again stratified by PID whereas (remote) homologs belonging to the same Pfam clan but different Pfam families, whose ground-truth PID is unknown, are all assigned to a special “Clans” bin. Unrelated pairs (negative class), whose ground-truth PID is also unknown, are included in all bins.

The results of this experiment are summarized in Figure 4. We also show in Supp. Figure S10 an alternative manner to evaluate performance, using the area under the precision-recall curve (AUPRC). We see that, unsurprisingly, all methods are largely successful in detecting homology for sequence pairs with moderate-to-large PID (≥ 0.2) in all settings. Interestingly, DEDAL is markedly better at detecting homology of highly divergent sequences across all four settings. For the smallest PID bin (< 0.1), DEDAL improves in-distribution performance by 25% (from AUROC=0.797 to AUROC=0.997) and 42% (from AUROC=0.696 to AUROC=0.992) for Pfam domains and extended domains, respectively. Once again, the performance of DEDAL degrades in the out-of-distribution split while remaining significantly superior to the baselines, boosting AUROC by 23% (from AUROC=0.770 to AUROC=0.948) and 30% (from AUROC=0.698 to AUROC=0.910) for domains and extended domains, respectively. When detecting homology for sequences belonging to the same Pfam clan but different Pfam families (“Clans” bin), the baselines perform only slightly better than random guessing. In the in-distribution split, their AUROC reaches 0.611 and 0.550 for Pfam domains and extended domains, respectively. In contrast, DEDAL’s performance in the in-distribution split for the “Clans” bin is comparable to that of homologs from the same Pfam family in the smallest PID bin (< 0.1), with corresponding AUROCs of 0.992 and 0.987 for domains and extended domains, respectively. In the out-of-distribution split, DEDAL exhibits a marked performance drop in the “Clans” bin compared to the PID< 0.1 bin, suggesting a certain degree of memorization of clan-specific motifs. Still, its performance remains well above that of the baselines, with AUROCs of 0.811 and 0.725 for Pfam domains and extended domains, respectively.

**Figure 4:**
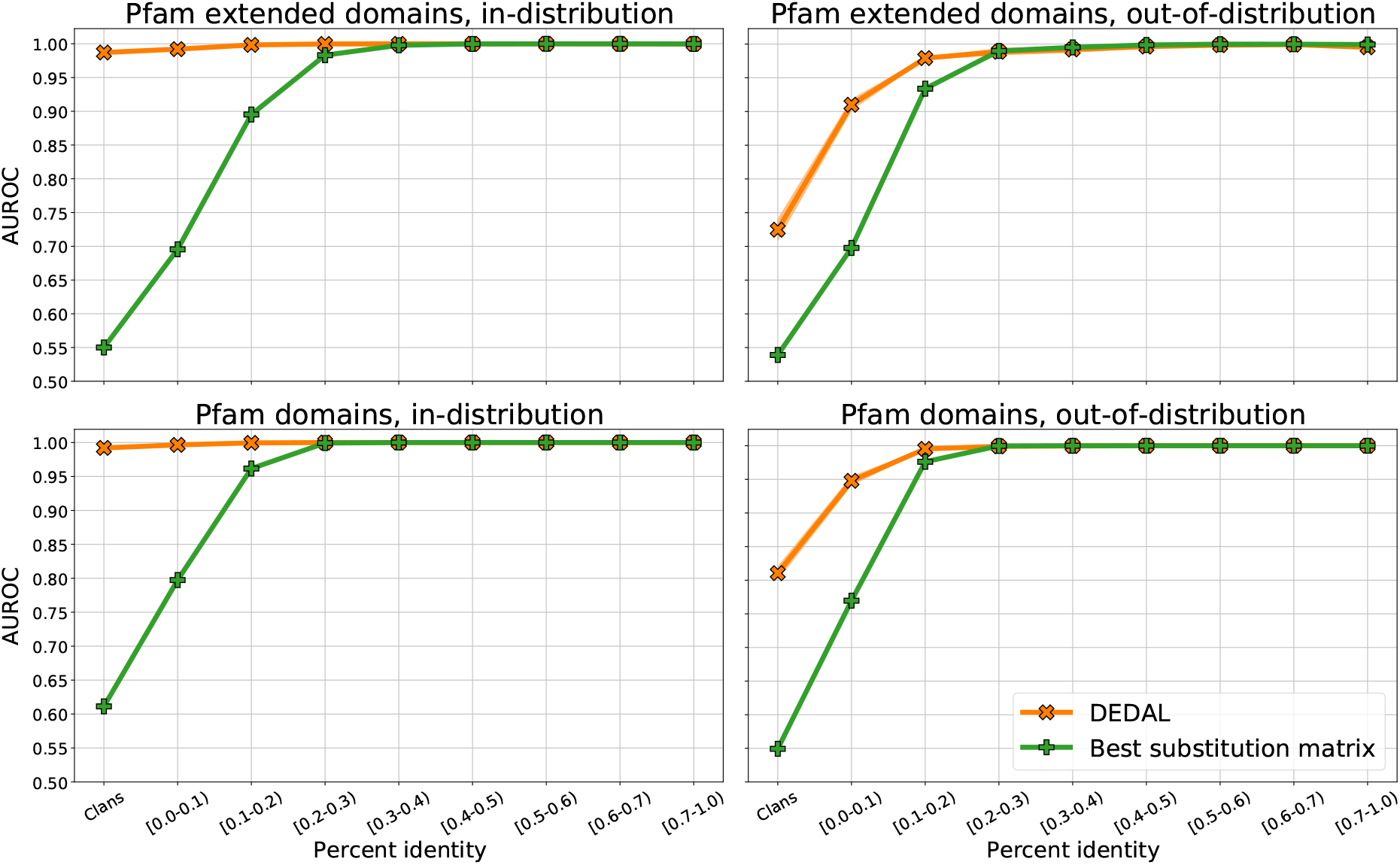
Homology detection AUROC of DEDAL and the best-performing substitution matrix baseline, in the in- and out-of-distribution settings (respectively, left and right columns), and for Pfam extended or raw domains (respectively, top and bottom rows).

## Discussion

Using recent advances in deep language models with transformers and novel differentiable alignment modules, we showed that DEDAL learns a continuous representation of protein sequences that, combined with the SW algorithm, leads to more accurate pairwise sequence alignment and homology detection than using the SW algorithm with fixed substitution matrices and gap penalties. The improvement is particularly striking in hard cases, where we want to align remote homologs with limited sequence identity, and we showed that the model learned by DEDAL on some sequence space generalizes well to new clans, suggesting that DEDAL learns universal biological properties not easily captured by standard substitution matrices and affine gap penalties.

Evaluating the performance of methods for remote homology detection is challenging, since by construction our current knowledge of homology relationships is mostly derived from existing bioinformatics tools, which have their own limitations. For the purpose of this study, we consider sequences in different Pfam clans (and with no nested domains) as non-homologous, which is only a proxy to the biological definition of homology. While there are arguably many unknown homologous pairs with very low sequence identity that we may be missing in current databases, we note that our definition of homology already creates many very challenging remote homology detection problems when sequences have low sequence identity, giving a useful benchmark to make progress on homology detection.

Since DEDAL incorporates a number of design choices, such as the architecture of the model or the strategies to train it, one may wonder which aspects are the most critical to explain its success. To address that question, we performed an ablation study where we systematically evaluated the effect of various design choices on the performance of DEDAL (Supplementary Results S2.1). In short, we found that replacing our rich parameterization of position-specific gap open and extend parameters by a simpler model where the gap open and extend parameters are position independent does not affect the performance of DEDAL much, achieving slightly superior alignment *F*_1_ scores in the in-distribution split but marginally inferior results for out-of-distribution sequence pairs. Another simplification would be to replace our position-specific affine gap penalty by a position-specific linear gap penalty, as for example used in DeepBLAST [25], which, however, we found leads to a small performance drop for remote homologs that is most pronounced in the out-of-distribution split.

On the technical side, we explored two approaches to create a differentiable SW alignment module, needed to train the parameters of DEDAL in the “learning to align” task, using either a smoothing technique [36] or a perturbation technique [37]; we found no significant difference between both in terms of performance, and implemented the one based on perturbations in the final DEDAL model. Regarding the set of alignments used to train DEDAL, we found that it is beneficial to use Pfam extended domains instead of Pfam domains when we want DEDAL to be able to predict accurate local alignments. Excluding sequences related to the out-of-distribution families from the “protein universe” seen when pretraining DEDAL on the masked language modeling task led to a slight performance drop for remote homologs, albeit insignificant relative to the performance gap with respect to the baselines. Regarding the strategy to train end-to-end jointly the transformer and parameterizer, we found that this is indeed significantly better than a more classical, two-step strategy that would first train a transformer encoder on the masked language modelling task, and second the parameterizer on the “learning to align” task by keeping the transformer fixed. This suggests that a universal language model, such as the one presented by [31], is not sufficient and should be at least fine-tuned for optimal performance in alignment. Finally, we assessed the benefit of context-dependent embeddings by simply training a model where the substitution cost is constrained to depend only on the amino acids to be aligned; unsurprisingly, we observed a strong drop in performance for this model, reaching about the same performance as the best-performing substitution matrix in the literature. All in all, our ablation study indicates that DEDAL is robust to changes in several technical aspects of the model, but that the main idea to jointly train a deep language model and a flexible parameterizer for SW is key to its performance.

This work opens several research directions for the future. First, as pure language models are increasingly used for a variety of downstream tasks, it is interesting to explore if adding alignment information to supervise language models, as DEDAL does, may also benefit other downstream tasks beyond sequence alignment. In preliminary experiments, we did not observe significant differences between the transformer encoder trained by DEDAL and one purely trained on a language modelling task on the TAPE benchmark [38], a collection of problems to assess the relevance of protein embedding models for various downstream tasks (see Supplementary Results S2.2), but believe there is room for improvement. Second, it will be interesting to extend DEDAL to the multiple alignment setting (MSA), either by combining pairwise alignment operations, as proposed by [26], which could directly benefit from DEDAL, or by directly embedding multiple non-aligned sequences and creating a differentiable MSA module. Third, given the accuracy of DEDAL’s alignment and ability to capture homology, it will be interesting to develop fast approximations based, e.g., on fast similarity search in continuous spaces, in order to allow accurate homology search and alignment in large databases.

## Methods

### Data

To train the transformer encoder with a masked language modelling task, we used 30, 162, 111 cluster representative sequences from the March 2018 release of the UniRef50 database [28]. These representatives provide a non-redundant yet complete coverage of UniProtKB [39], obtained by clustering sequences with at least 50% percent identity and 80% overlap with the cluster’s longest sequence using MMseqs2 [40]. A representative for each cluster was chosen accounting for factors such as quality of the UniProtKB entry and availability of annotations, among others. Given the relatively low redundancy between UniRef50 representative sequences, sequences were allocated to train or test uniformly at random as in [31], resulting in training and test sets of size 27,052,653 and 3,019,006, respectively. We used the training set to define the loss of the masked language model task, and the test set to monitor its convergence during optimization of the training loss. A small validation set of 90,452 sequences was further reserved for hyperparameter tuning. For computational efficiency, we limit the length of sequences fed to the model for this task to at most 1,023 amino acids. Longer sequences are cropped to length 1, 023 with a uniformly distributed offset sampled on-the-fly at training and evaluation time.

To create pairs of homologous sequences with known alignment, we collected 19, 179 manually curated multiple sequence alignments (MSAs) from release 34.0 of the Pfam-A seed database [29], grouped into 12, 452 evolutionarily-related clans (including 645 Pfam clans that altogether comprise 7,372 families, and the remaining 11,807 families that we assigned to their own clan). Together these span a total of 1,223,021 sequences from which we kept only the 1, 198, 870 sequences (98% of the total) of length smaller than 512. We define a pair of homologous sequences as a pair of sequences in the same clan, from which we extract a correct pairwise alignment when both sequences are in the same family. Any two sequences not in the same clan are deemed as not homologous in the homology detection task if they share no other annotations at the clan level. Sequences not in the same clan but with common annotations (as could be the case, e.g., for nested domains) are deemed ambiguous and excluded from the analysis^1^. From this set of Pfam domain sequence pairs, we also build a set of Pfam extended domain sequence pairs by adding flanking sequences with random lengths upstream and downstream of each domain sequence. For each pair of sequences, these flanks are randomly chosen (sub)sequences from the UniProtKB protein sequence database sharing no clan annotations with neither the domain sequence nor flanks of the other Pfam extended domain sequence in the pair (see Supplementary Methods S1.1). As a result, the correct alignment between two homologous extended domains usually does not span the full sequences. For both Pfam domains and extended domains, we split the full dataset into five disjoint splits: train (80%), in-distribution validation (2.5%), out-of-distribution validation (2.5%), in-distribution test (7.5%) and out-of-distribution test (7.5%). To this end, we first select 255 and 955 out of the 12, 452 Pfam clans uniformly at random to form the out-of-distribution validation and out-of-distribution test, respectively. The number of clans was chosen so that these splits contain approximately 2.5% and 7.5% of the total number of sequences. Finally, we divide the set of sequences in the remaining 10, 493 clans uniformly at random to form the train, in-distribution validation and in-distribution test splits, so that a single family may have sequences in all three sets.

The alignment task is then defined by *all* pairs of sequences in each split belonging to the same Pfam family, resulting in 126,176, 858 training, 111, 031 in-distribution validation, 1, 101, 061 in-distribution test, 5, 059, 395 out-of-distribution validation and 11, 558, 433 out-of-distribution test samples, respectively. For the homology task, we complement these homologous pairs (positive class) with pairs of sequences belonging to different Pfam clans (negative class) as well as pairs of sequences belonging to the same Pfam clan but different Pfam families (“hard” cases in the positive class). These are sampled uniformly at random from their respective sets so that they make 50% and 11% of the total data in each split, respectively. These proportions were motivated by having a balanced class ratio as well as a moderate amount of remote homologs in the positive class (roughly 25% including sequence pairs from the same Pfam family with PID < 0.1).

### DEDAL model

DEDAL is a parametric model that transforms a pair of sequences onto three matrices of gap open, gap extend penalties and substitution costs, respectively, and then uses the SW algorithm to align the two initial sequences using these matrices of parameters. Each protein sequence is seen as a sequence of tokens in a 27-letter alphabet (22 amino acids including pyrrolysine (O) and selenocysteine (U), 3 ambiguous codes B, Z and X, and 2 special tokens `<EOS>`, which we append to each sequence, and `<MASK>`, which is used for masked language modelling only). Each such sequence is mapped to a sequence of vectors by a transformer encoder neural network similar to the smallest transformer-based model of [31] which uses *L* = 6 transformer encoder layers with *h* = 12 heads per layer and embeddings of dimension *d* = 768. Each transformer encoder layer combines multi-head self-attention, position-wise multilayer perceptron, dropout and layer normalization. From the output of the transformer, we compute the substitution score between two positions as a bilinear function of their vector representations, and the gap and open penalties as the softplus function applied to a bilinear function of their vector representations to ensure they are nonnegative. An in-depth description of the DEDAL model as well as of its architecture and hyperparameters can be found in Supplementary Methods S1.2.

### Differentiable SW alignment

Given two sequences *x* = *x*_1_ … *x_n_* and *y* = *y*_1_ … *y_m_* of amino acids, an alignment *π* is a subset of distinct positions in *x* and *y* that are matched in increasing order. Let us denote by Π(*x, y*) the set of possible alignments between *x* and *y*, including the empty one. Given three *n* × *m* matrices *S* (substitution scores), *O* (gap open) and *E* (gap extend penalties), the local alignment score *s*(*π*) of a candidate alignment *π* is defined as the sum of the substitution scores over matched positions, and gap penalties defined as the gap open penalty where a gap starts and gap extend penalties where it continues. The SW algorithm finds the maximum scoring alignment with the dynamic programming recursion over the (*n* + 1) × (*m* + 1) matrices *M*, *X* and *Y* defined for any *i* = 1, …, *n* and *j* = 1, …, *m* by *M*_*i*,0_ = *M*_0,*j*_ = *X*_*i*,0_ = *X*_0,*j*_ = *Y*_*i*,0_ = *Y*_0,*j*_ = −∞ and:

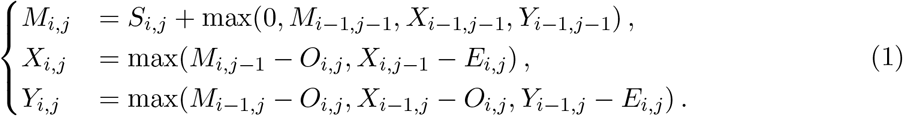

The score of the best alignment is then 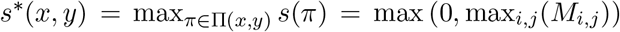, and the best-scoring alignment 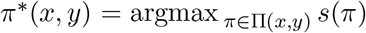 can be recovered by backtracking the argmax in the recursion. Both operations can be implemented in *O*(*nm*) complexity in time and space; to benefit from modern computational infrastructure, we implemented a version optimized for hardware accelerators such as GPU and TPU, which practically runs in *O*(max(*m, n*)) time and in parallel across a batch of sequences to align, using a wavefront algorithm (see Supplementary Methods S1.3). We note that the score of any alignment *π* ∈ Π(*x, y*) is a linear function of the parameters Θ = (*S, O, E*) seen as a 3*nm*-dimensional vector, which we write as *s*(*π*) = Θ^⊤^Φ(π) for some mapping 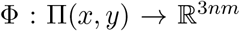. This shows that *s*\*(*x, y*) is almost everywhere differentiable in Θ, with gradient Φ(*π*\*(*x, y*)), which potentially allows to backpropagate to Θ any differentiable objective function that depends on the optimal alignment score, such as the one used in the homology detection task. On the other hand, the best alignment *π*\*(*x, y*) is a piecewise constant function of Θ, and any loss that only depends on *π*\*(*x, y*), such as how different it is from the correct alignment, can not be backpropagated to the parameters. In the “learning to align” task, however, we need to have a differentiable loss to minimize with respect to Θ when SW finds an alignment *π*\*(*x, y*) but the correct alignment is 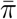, potentially different from *π*\*(*x, y*).

To overcome this issue, a first idea is to consider the so-called structured perceptron loss [41], which penalizes the difference between the score of the best alignment and the score of the true alignment: 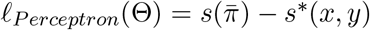. Its gradient is simply 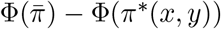, which can be computed with the SW forward and backward recursions. To create a smoother loss function, we consider two natural extensions to the perceptron loss. The first one is the so-called CRF loss [42] given by 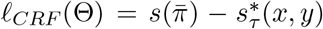 for *τ* ≥ 0. Instead of being the maximum score, 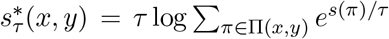 is the soft-maximum score. The CRF loss can also be interpreted as a negative log-likelihood model under a Gibbs distribution, and as shown in [36], its value and gradient can be computed as efficiently as for the perceptron loss, by simply replacing the max operation in the SW recursion (1) by a softmax of the form 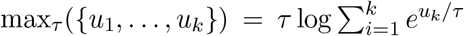, and adapting the backtracking. The CRF loss thus provides a smooth approximation to the perceptron loss, which converges to it when *τ* goes to zero.

The second extension creates smoothness by random perturbation of Θ, as proposed by [37]. More precisely, we consider a random perturbation Θ + *τZ* of the parameters Θ, where *τ* ≥ 0 and *Z* is a standard multivariate normal random vector; then the so-called Fenchel-Young loss [43] associated to the perturbation according to [37] is a valid loss with gradient 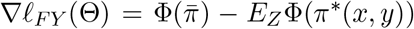. In other words, *ℓ_FY_* is a valid loss whose gradient is the expectation of the perceptron loss gradient under random perturbation of the SW parameters. In practice we compute a stochastic gradient (which is enough to optimize the loss function by stochastic gradient descent) by taking the perceptron loss gradient after a single random perturbation of the parameters by *τZ*. We note, again, that the Fenchel-Young loss is a smooth approximation to the perceptron loss, which converges to it when *τ* goes to 0.

Importantly, both extensions not only encourage the correct alignment to have a “large” score, but they also encourage it to have a “better” score than other candidate alignments. Intuitively, one can see this by noticing that the CRF loss, for example, is the negative log-probability of the correct alignment among all possible alignments under a Gibbs distribution where the probability of each alignment *π* is proportional to *e^s(π)/τ^*; since the probabilities of all alignments sum to 1, encouraging the probability of the correct alignment to increase (by minimizing the CRF loss) can only be achieved by not only increasing the score of the correct alignment, but also decreasing the scores of other, alternative alignments, hence encouraging the correct alignment to be the unique maximizer of the alignment score.

#### Homology detection classifier

We treat homology detection as a binary classification problem, aiming to elucidate whether a pair of sequences *x* and *y* belong to the same Pfam family or not, given their alignment score *s*\*(*x, y*). Since it is well known that the distribution of alignment scores on random sequences strongly depends on their length [44], we must correct for the length effect on the score and pose the following model, inspired by how E-values are defined to assess the significance of alignment scores, for the probability *P* that *x* and *y* are related given their alignment score and lengths *n* and *m*:

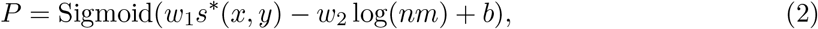

where *w*_1_, 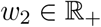 and 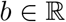 are parameters. These parameters are jointly trained end-to-end alongside the sequence encoder and the differentiable alignment layer using a binary crossentropy loss. The differentiable alignment layer is thus explicitly trained to output small scores for unrelated sequence pairs, preventing the model from acquiring biases caused by it only being trained on related sequences by the alignment task.

#### Output head for masked language modelling

We follow [30, 31] and define a masked language modelling task to train the transformer by randomly selecting 15% of the residues in each sequence to be prediction targets. 80% of these are substituted by a special `<MASK>` token, 10% by a randomly chosen residue and the remaining 10% is left unchanged. For a sequence *x* of length *n*, we denote by 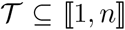 the set of target residues, chosen uniformly at random, and by 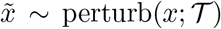 the resulting randomly perturbed sequence. The goal of the masked language modelling output head is to predict the original value of each perturbed token *x_i_* from the embeddings of the (perturbed) sequence 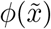 as

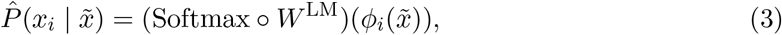

where 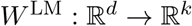 is a learnable, affine mapping that is *not* tied to *F* in the input embedding layer (see Supplementary Methods S1.2.1). These predicted probabilities 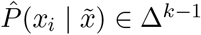 are then compared to the ground-truth using a cross-entropy loss,

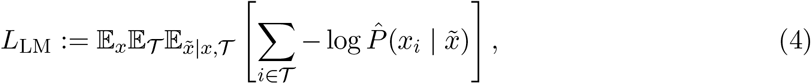

where we omitted the dependence of *L*_LM_ on the parameters *ϕ* of the sequence encoder and *W*^LM^.

#### Training DEDAL

We first pretrain the sequence encoder on the masked language modelling task for 1 million steps. After that, we jointly train the sequence encoder, differentiable alignment layer, homology detection classifier and masked language modelling output head for 1.3 million additional steps to minimize a multi-task objective:

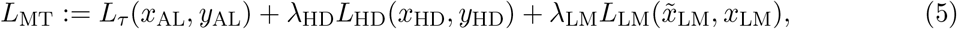

where *L_τ_* is a loss for the alignment task dependent on the relaxation scheme, *L*_HD_ the binary cross-entropy loss for the homology detection task, *L*_LM_ the cross-entropy loss for the masked language modelling task and 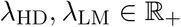 are hyperparameters controlling the relative weight of each loss on the multi-task objective. The batches (*x*_AL_, *y*_AL_), (*x*_HD_, *y*_HD_) and 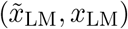 for the alignment, homology and masked language modeling tasks are sampled independently of each other at each training step. We found it advantageous to disable the homology task (effectively using λ_HD_ = 0) for the first 300, 000 steps. In all three training phases (pretraining, fine-tuning without homology and fine-tuning with homology), we use the Adam [45] optimizer with a linear warm-up of 8, 000 steps until reaching a maximum learning rate lr_max_ that we treat as a hyperparameter, followed by an inverse square root decay schedule.

### Metrics

Given two homologous sequences *x* and *y* from an evaluation set 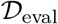 with known correct alignment, let us denote by *m*_true_(*x, y*) the set of pairs of indices aligned to each other in the correct alignment, and by *m*\*(*x, y*) the set of pairs of indices predicted to be aligned by a SW prediction. We assess the performance of the SW prediction in terms of precision, recall and *F*_1_ score defined as:

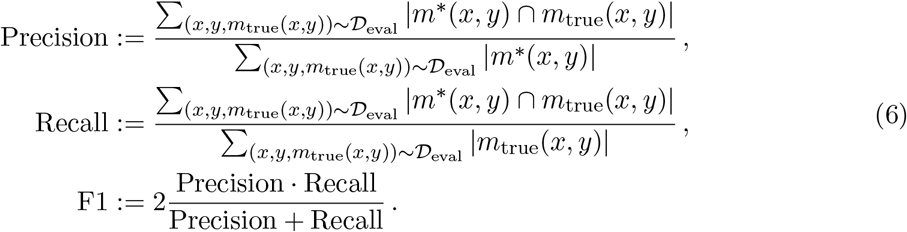

To evaluate homology detection performance, we use the Area Under the ROC Curve (AU-ROC) and the Area Under the Precision-Recall Curve (AUPRC).

To better reflect the relative difficulty of each task, we stratify all alignment and homology detection metrics by Percent IDentity (PID) [46], using the definition of PID from [47], namely,

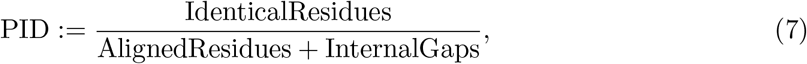

where the number of identical residues and internal gaps is computed for each pair of related sequences using the ground-truth alignments. For homology detection, only a subset of the positive class (sequence pairs belonging to the same Pfam family) can be stratified in this manner. Negative pairs, for which no ground-truth alignments are available, are shared across all PID bins while positive pairs belonging to different Pfam families, whose curated PIDs are also unknown, are stratified as their own “Clans” bin. Since we chose a version of PID that disregards external gaps, it is largely unaffected by our data augmentation process where we add flanking regions to Pfam domains (Supplementary Figure S7).

All reported metrics were evaluated in the entirety of the (in- and out-of-distribution) test splits for both the Pfam extended and raw domain settings.

### Baselines

We compared the performance of DEDAL on the alignment and homology detection tasks to three widely-used families of substitution matrices: BLOSUM [33], VTML [34, 35] and PFASUM [20]. For each family, we exhaustively optimized over the following hyperparameters: 1) the matrix number in the family, where we considered four matrix numbers for the BLOSUM family (45, 50, 62, 80), six for the VTML family (10, 20, 40, 80, 160, 200) and, finally, seven for the PFASUM family (31, 43, 51, 60, 70, 80, 90); 2) the gap open penalty, taking any integer value between 5 and 18; 3) the gap extent penalty, taking values in {0.5,0.75,1.0,1.25,1.5,2.0}. We selected the best-performing configuration over these 1,428 combinations as the baseline representing substitution matrices, following the protocol described in Supplementary Methods S1.5.

## Data availability

The Uniref50 dataset is freely available under the Creative Commons Attribution (CC BY 4.0) License from https://www.uniprot.org. Pfam is freely available under the Creative Commons Zero (“CC0”) licence from https://pfam.xfam.org

## Code availability

The code used in this study is freely available under an Apache 2.0 license at https://github.com/google-research/google-research/tree/master/dedal

## Author contributions

F.L.L. contributed to the development of the method, implemented most of the code, designed and ran most experiments, drafted the manuscript. Q.B. contributed to the development of the method, to the code, and to the experiments. M.B. contributed to the development of the method and to the code. O.T. designed and implemented major parts of the code and contributed to the experiments. J.P.V. designed the initial project, contributed to the method development and drafted the manuscript. All authors provided regular feedback to all aspects of the work, reviewed code and contributed to the writing of the final manuscript.

## Supplementary Material

### S1 Supplementary Methods

#### S1.1 Pfam domain sequence extension

As explained in the main Methods, we use pairs of aligned domain sequences from Pfam-A seed as ground-truth alignments, to supervise the “learning to align” task of DEDAL and assess the performance of various alignment algorithms. A drawback of using Pfam-A seed alignments as targets, however, stems from the fact that these are *de facto* global alignments. This is illustrated in Figure S1a, where we see that the vast majority of pairwise alignments from the original Pfam-A seed dataset have no flanking sequence tails outside the alignment region. Thus, models naively trained using these alignments as supervision targets will be heavily biased towards global alignment, which might impede their ability to predict local alignments whenever necessary (c.f. “Trained on Pfam domains” ablation in Figures S5 and S6).

**Figure S1:**
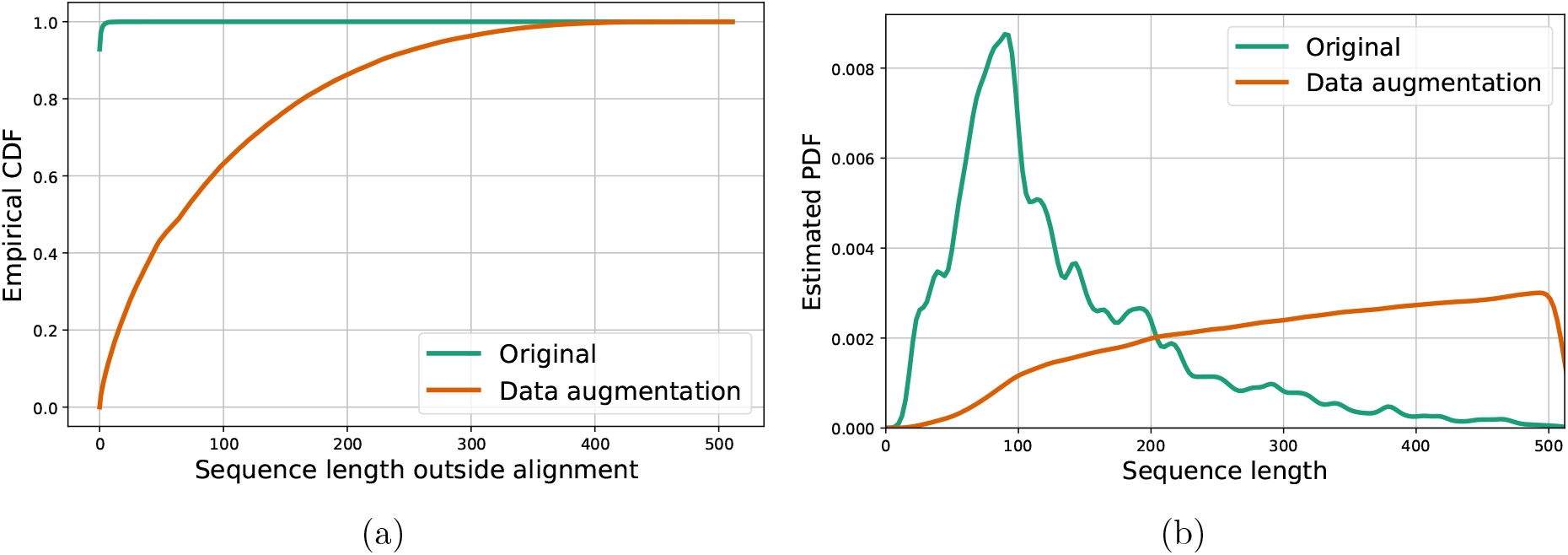
(a) The empirical CDF of the number of residues not mapping to matches or internal gaps in a target alignment. (b) A kernel density estimate of the sequence length distribution. A random set of 1, 048, 576 sequence pairs from the training set was used for both figures.

To overcome this limitation, we apply *data augmentation* to the Pfam-A seed dataset, creating *local* pairwise sequence alignment targets by adding flanking sequences to the original Pfam domains. To this end, every time a Pfam-A seed sequence is drawn as part of a sequence pair for training or evaluation, we first sample lengths for the N-terminus and C-terminus flanks to be added as

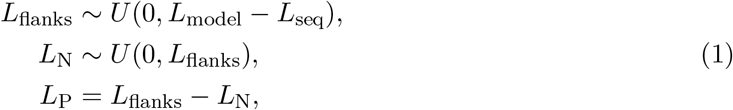

with *L*_seq_ being the length of the original Pfam-A seed domain sequence and *L*_model_ = 511 the maximum sequence length we consider for this task. In other words, the total length of the flanking sequences being added is sampled uniformly between zero and the maximum possible, given the computational budget. These *L*_flanks_ residues are subsequently allocated to N and C-terminus uniformly at random among all valid choices. Next, for each of the two flanks, we sample a random (sub)sequence of length *L*_N_ (resp. *L*_P_) from UniProtKB. To prevent the inclusion of flanks from causing any test set “leakage”, we ensure that:

1. UniProtKB sequences that match any of the Pfam families in the out-of-distribution validation or out-of-distribution test splits may *not* be used to generate flanks.
2. The same UniProtKB sequence may *not* be used to generate flanks for Pfam-A seed sequences in different splits.
3. UniProtKB sequences that contain Pfam-A seed domain sequences may only be used to generate flanks for Pfam-A seed sequences in the same split to which those sequences were allocated^1^.

Furthermore, we aim to minimize the number of erroneous “ground-truth” extended alignments by guaranteeing that, for every pair of extended Pfam domain sequences, the flanks added to one sequence have no known clan annotations in common with neither the Pfam-A seed region nor the flanks of the sequence^2^. To further increase the data diversity, the N and C-terminus flank lengths as well as the flanking sequences themselves are sampled independently for each sequence pair. In other words, the same Pfam-A seed domain sequence may be extended differently depending on the other sequence it is paired with.

As shown in Figure S1a, this data augmentation process greatly increases the diversity of alignment targets relative to the original data. Moreover, the augmented input sequences have a significantly less skewed length distribution, as can be seen in Figure S1b.

The correctness of the proposed data augmentation scheme is partly limited by the coverage of existing clan-level annotations. Because of this, it is possible that a pair of Pfam-A seed domain sequences is extended with flanks that are related by a not-yet-known homology relationship. To corroborate that such occurrences are not prevalent enough to affect our results in any significant manner, we also experimented with an alternative data augmentation scheme using *synthetic* flanks. These are simply obtained by sampling residues i.i.d. using the background amino acid frequencies estimated from Pfam-A seed sequences. The resulting Pfam synthetic domains are arguably less realistic, as well as challenging for DEDAL, which was trained specifically to align “protein-like” sequences. In spite of this, Figures S2 and S3 show that the performance of DEDAL degrades only slightly when evaluating on sequences with synthetic flanks, despite having been trained only on “protein-like” sequences with UniProt-based flanks. Moreover, the baselines perform similarly with both types of flanking sequences, suggesting that erroneous ground-truth alignments due to missing annotations, while possible, are rare enough not to compromise our evaluation.

**Figure S2:**
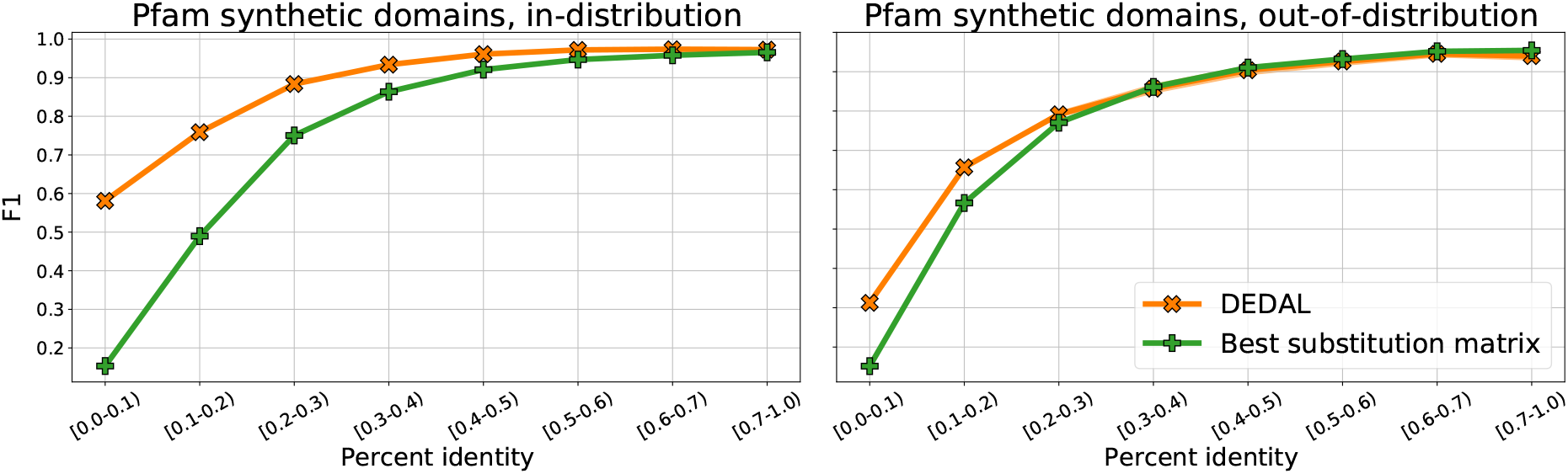
Alignment *F*_1_ score of DEDAL (trained on UniProt-based flanks) and the best-performing substitution matrix baseline, in the in- and out-of-distribution settings (respectively, left and right columns) for Pfam domains extended with synthetically-generated flanks (sampled i.i.d. from the background amino acid-frequencies in Pfam-A seed).

**Figure S3:**
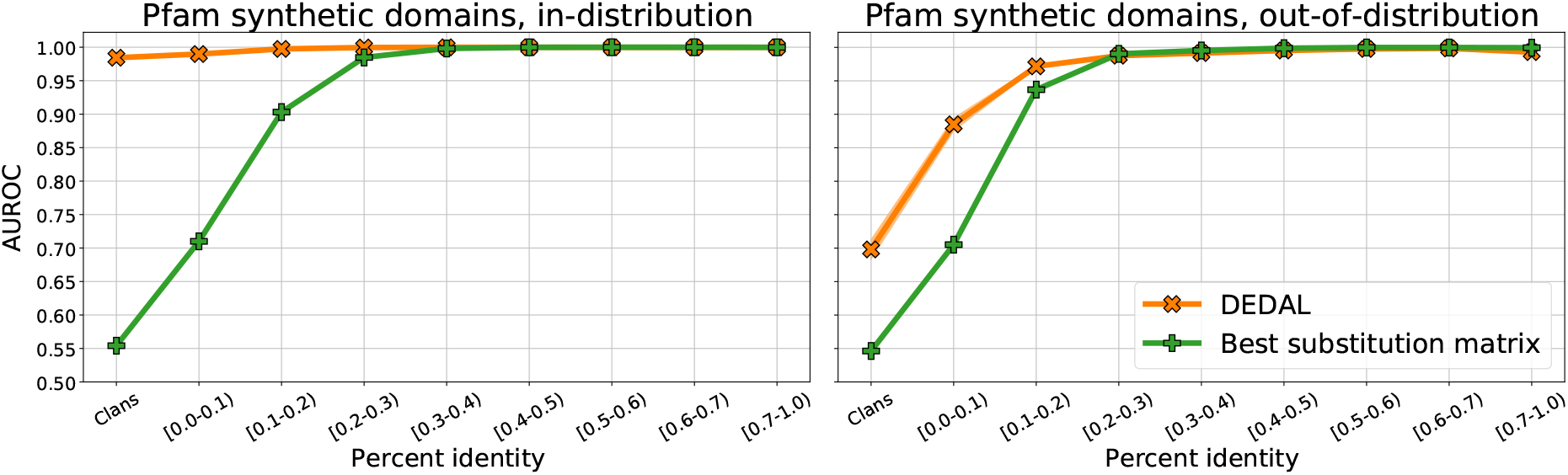
Homology detection AUROC of DEDAL (trained on UniProt-based flanks) and the best-performing substitution matrix baseline, in the in- and out-of-distribution settings (respectively, left and right columns) for Pfam domains extended with synthetically-generated flanks (sampled i.i.d. from the background amino acid-frequencies in Pfam-A seed).

#### S1.2 DEDAL model and training

In this section we provide additional details on the DEDAL model, and how it is trained. DEDAL contains a *sequence encoder* that computes continuous representations of protein sequences and multiple task-specific *output heads* that make predictions based on these continuous representations. All model components are jointly trained end-to-end using a multi-task loss, with the exception of the output heads for downstream tasks, which are fine-tuned independently for each task. The remainder of this section introduces each model component in turn, concluding with a description of the training process.

##### S1.2.1 Contextual sequence embeddings

The sequence encoder is a parametric function that maps sequences *x* = (*x*_1_, *x*_2_, …, *x_n_*) over a finite alphabet (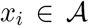 for all *i* = 1, …, *n*) to a sequence of *d*-dimensional vectors of the same length 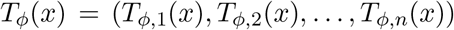. Crucially, its architecture was chosen so that the *embedding* for the *i*-th input residue 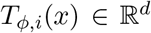 depends on the entire sequence *x* rather than on *x_i_* alone, making these sequence embeddings *contextual*. To this end, we use a transformer encoder [1]. Transformer-based models achieve state-of-the-art performance across a wide range of problems in natural language processing and, more recently, computer vision. Unlike convolutional or recurrent architectures, transformers make extensive use of *attention* to model long-range interactions between tokens directly, thus excelling at capturing non-local context.

###### Tokenization

In our case, the alphabet 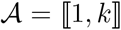 consists of *k* = 27 *tokens* corresponding to (i) the 20 standard amino acids, (ii) the less common Pyrrolysine (O) and Selenocysteine (U) amino acids, (iii) the “ambiguous” amino acid codes B, Z and X and (iv) the special tokens `<EOS>`, which we append to each sequence, and `<MASK>`, which is used for masked language modelling only.

###### Input embeddings

Similar to [1, 2], the first layer of our sequence encoder is an *input embedding layer* that additively encodes residue identity and position information for each token in the input sequence. Given a sequence 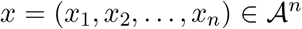, we define 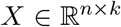 to be the *one-hot* representation of *x*, i.e., *X_i,j_* = *δ_j_* (*x_i_*). The input embedding layer computes non-contextual, continuous representations for each residue as

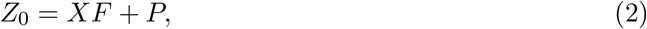

where 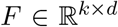 contains learnable input embeddings for each token in 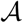 and 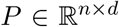 is a matrix of sinusoidal, absolute position embeddings [1].

###### Transformer encoder layers

The non-contextual embeddings 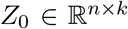 are then fed to a stack of *L transformer encoder layers* which use *multi-headed self-attention* followed by a position-wise multilayer perceptron (*MLP*). Together, these adaptively incorporate into each residue’s embedding information about the other tokens in the sequence. In particular, we use transformer encoder layers with pre-layer normalization [3, 4],

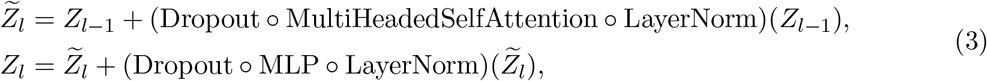

where ∘ denotes function composition.

The multi-headed self-attention mechanism combines the output from *h* independent self-attention layers as

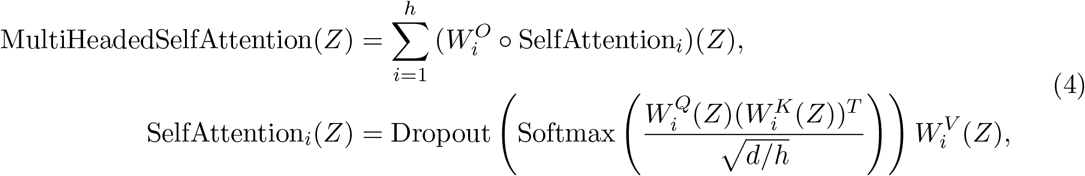

where 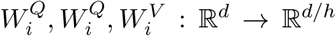 and 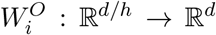 are all learnable, affine mappings. These and the softmax non-linearity are applied row-wise to their respective *n* × *d, n* × *d/h* and *n* × *n* inputs.

The MLP contains a single hidden layer that uses a Gaussian Error Linear Unit (GELU) [5] element-wise non-linearity, namely,

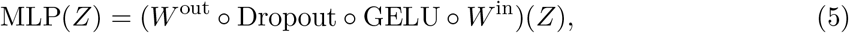

where 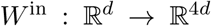 and 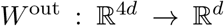 are also learnable, affine mappings applied row-wise to their inputs.

###### Output normalization

Finally, layer normalization is applied to the output of the *L*-th transformer encoder layer to obtain the contextual embeddings 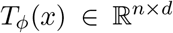 for an input sequence 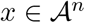,

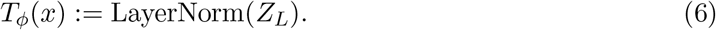

##### S1.2.2 Parameterizer

Once each sequence is mapped to a continuous representation, we compute matrices of parameters to be used by the SW alignment algorithm from the continuous representations. This allows the SW algorithm to rely on information other than the identity of individual residue pairs by modelling the substitution scores, gap open and gap extend penalties as a parametric function of the (contextual) embeddings *T_ϕ_*(*x*) and *T_ϕ_*(*y*) of the sequences to be aligned. In practice we implement these as symmetric bilinear forms,

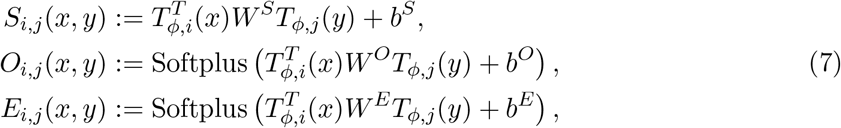

where 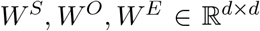 are symmetric matrices and 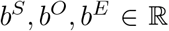 scalar biases, respectively. Together, these form the set of parameters *β* of the parameterizer function *P_β_*. An element-wise softplus non-linearity is used to ensure gap penalties remain non-negative.

This model is a strict generalization of the traditional parameterization of the SW algorithm based on substitution matrices. We can recover these as a particular case by (i) using a one-hot representation as embeddings, i.e., *T_ϕ_*(*x*) = *X* and *T_ϕ_*(*y*) = *Y*, (ii) restricting the gap penalties to be position-independent, i.e., *O_i,j_* ≔ *O, E_i,j_* ≔ *E* with 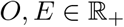 and (iii) fixing 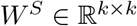 to the desired substitution matrix. In contrast, we jointly train the transformer-based sequence encoder and the alignment layer, resulting in a substantially more expressive parameterization.

##### S1.2.3 Architecture and hyperparameters

We adopt the architecture of the smallest transformer-based model in [2], which uses *L* = 6 transformer encoder layers with *h* = 12 heads per layer and embeddings of dimension *d* = 768. Glorot initialization [6] is used for all weight matrices with the sole exception of the input embeddings table, which are initialized from a standard normal distribution. The dropout rate is set to 0.1 throughout. In all experiments, a global batch size of 128 sequences (resp. sequence pairs) was used for the masked language modelling and alignment tasks while the homology task used 256 sequence pairs. No attempts were made to optimize over these hyperparameters. Additionally, DEDAL was trained with maximum learning rate lr_max_ = 10^−4^ and multi-task loss weights λ_HD_ = λ_LM_ = 20. Unless stated otherwise (e.g. ablations), DEDAL used the perturbation-based approach in [7] with relaxation strength *τ* = 0.1 to smooth the SW alignments during training. For computational considerations, these hyperparameters were selected after a lightweight exploration of model performance on the in-distribution and out-of-distribution validation splits, which have no overlap with the held-out test set.

#### S1.3 Accelerator-friendly implementation of the (smooth) SW algorithm

As explained in the Methods section, we use the SW algorithm to find the maximum scoring alignment (or a soft maximum in the case of the CRF loss) with the dynamic programming recursion over the (*n* + 1) × (*m* + 1) matrices *M*, *X* and *Y* defined for any *i* = 1, …, *n* and *j* = 1, …, *m* by *M*_*i*,0_ = *M*_0,*j*_ = *X*_*i*,0_ = *X*_0,*j*_ = *Y*_*i*,0_ = *Y*_0,*j*_ = −∞ and:

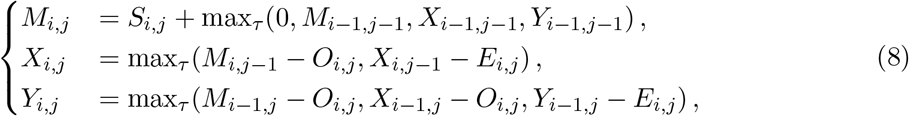

where 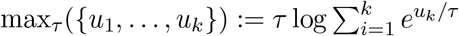 is the soft-max when *τ* > 0, or the max operator when *τ* = 0. The (soft) best-scoring alignment needed to compute the gradient of the score and of the different alignment loss functions with respect to *S*, *O* and *E* is then computed by appropriate backtracking.

The widespread availability of modern hardware accelerators such as Graphical Processing Units (GPUs) and Tensor Processing Units (TPUs) has been partly responsible for the recent success of deep learning models. These hardware resources excel at vectorized computation, perhaps most notably matrix-matrix and matrix-vector multiplication, which are the fundamental building blocks of most commonly-used deep learning layers. In contrast, the dynamic program underlying the SW algorithm appears to be inherently sequential. Arguably, its most natural description involves firstly iteratively computing *M_i,j_*, *X_i,j_* and *Y_i,j_* using a double for loop along the lengths of sequences *x* and *y*, followed by backtracking. Unfortunately, such an implementation would only profit from vectorization along the batch axis, i.e., carrying out the dynamic program updates for multiple sequence pairs simultaneously, remaining otherwise purely sequential along both length axes *i* and *j*. To prevent the differentiable alignment layer from becoming a computational bottleneck, we implement a wavefront transformation that allows expressing the SW algorithm in terms of a single `for` loop, leading to a computational footprint comparable to that of Recurrent Neural Networks (RNNs). Most importantly, we aim to accomplish this using solely operations supported off-the-shelf by frameworks such as TensorFlow [8], JAX [9] or PyTorch [10] for maximum compatibility and ease of use.

**Table S1:** Effect of applying the mapping *f*^(+)^ (resp. *f*^(−)^) to a tensor 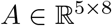, representing SW parameters for a pair of sequences of lengths *n* = 5 and *m* = 9. Entries in the output tensor not corresponding to any entry of *A* are padded with +∞ (resp. −∞).

|  |  |  |  |  |  |  |  |  |  |  |  |
| --- | --- | --- | --- | --- | --- | --- | --- | --- | --- | --- | --- |
| 1,1 | 1,2 | 1,3 | 1,4 | 1,5 | 1,6 | 1,7 | 1,8 | $\pm\infty$ | $\pm\infty$ | $\pm\infty$ | $\pm\infty$ |
| $\pm\infty$ | 2,1 | 2,2 | 2,3 | 2,4 | 2,5 | 2,6 | 2,7 | 2,8 | $\pm\infty$ | $\pm\infty$ | $\pm\infty$ |
| $\pm\infty$ | $\pm\infty$ | 3,1 | 3,2 | 3,3 | 3,4 | 3,5 | 3,6 | 3,7 | 3,8 | $\pm\infty$ | $\pm\infty$ |
| $\pm\infty$ | $\pm\infty$ | $\pm\infty$ | 4,1 | 4,2 | 4,3 | 4,4 | 4,5 | 4,6 | 4,7 | 4,8 | $\pm\infty$ |
| $\pm\infty$ | $\pm\infty$ | $\pm\infty$ | $\pm\infty$ | 5,1 | 5,2 | 5,3 | 5,4 | 5,5 | 5,6 | 5,7 | 5,8 |

Suppose 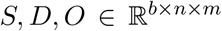 are tensors representing the (contextual) substitution scores, gap open and gap extend penalties of a batch of *b* sequence pairs of lengths at most *n* and *m*, respectively. Entries corresponding to sequences strictly shorter than *n* (resp. *m*) are assumed to have been padded with −∞ for S and +∞ for *D* and *E*. We also assume w.l.o.g. that *n* ≤ *m*. In order to vectorize the SW algorithm, we exploit that the dynamic program updates can be computed in parallel for all entries (*i, j*) along the same anti-diagonal, i.e., *i* + *j* = *k* for some 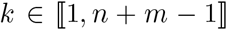. To reduce the number of non-contiguous indexing operations, we define mappings 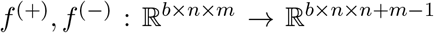 to rearrange the inputs *S, D* and *E* mapping their anti-diagonals to a single axis in the output tensors. Namely,

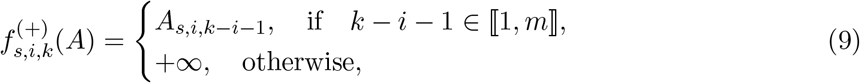

with *f*^(−)^ defined analogously but using −∞ as a sentinel value for invalid entries. A conceptual illustration of this transformation is shown in Figure S1. Setting 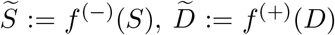 and 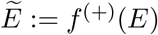, we can rewrite the dynamic program updates in Equation 8 as

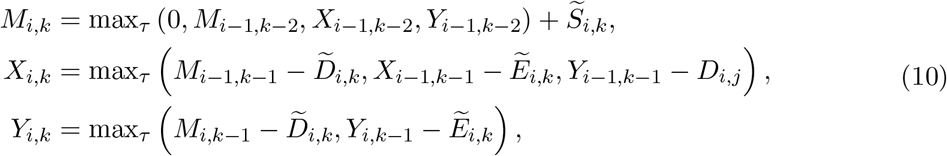

which can be trivially vectorized along the batch and *i* axes, requiring only a single `for` loop on 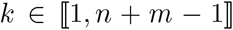. Despite the fact that this change does not alter the computational complexity of the algorithm, we found reducing the amount of strictly sequential computation to have a dramatic impact on runtime in practice, as shown in Table S4.

#### S1.4 Efficiently differentiating the soft-maximum SW score

The gradient of the soft-maximum score 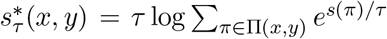 can in principle be computed using modern automatic differentiation (AD) software [8–10]. However, we have found using a custom implementation of 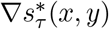 advantageous occasionally, particularly in terms of memory usage. For the sake of completeness, we include a detailed derivation of this custom implementation in this section.

Suppose that the recursive updates described in Equation 8 have been run until termination and define 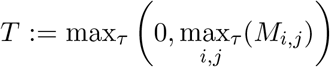 such that 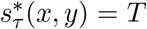. Our derivation will proceed in two steps. First, we express the sought-after partial derivatives 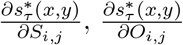 and 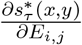 as a function of the *adjoint operators* 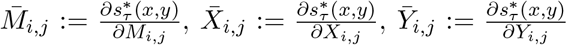 and of the (local) partial derivatives of the update rules in Equation 8. Next, we derive update rules to iteratively evaluate all of these adjoint operators.

**Figure S4:**
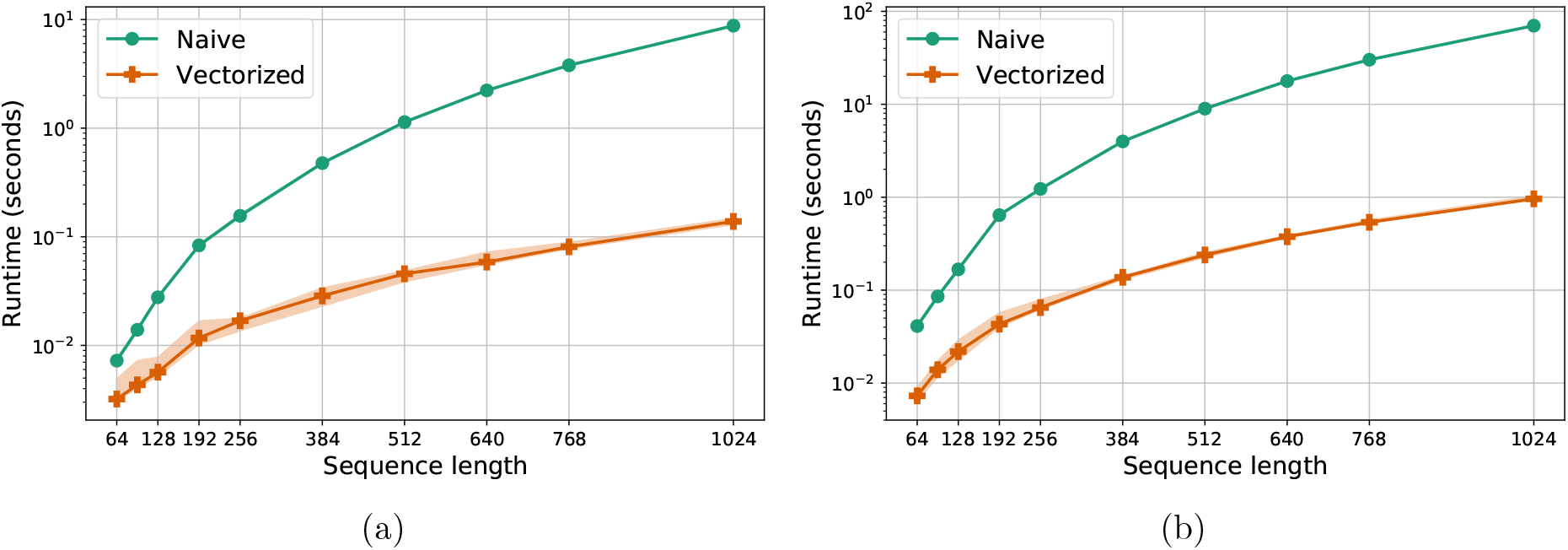
Runtime to align a batch of sequence pairs as a function of the length of the sequences for the proposed vectorization approach and a naive baseline implemented as a double for loop. Times shown reflect the forward pass only, computed in inference mode (*τ* = 0) using 4 TPU v3 chips. All data points were obtained as the median of 10 trials, with error bars reflecting the minimum and maximum runtimes among these. (a) Batch size fixed to 128 sequence pairs. (b) Batch size fixed to 1024 sequence pairs.

Using the chain rule, we may write

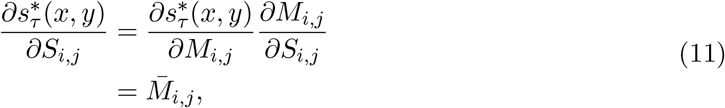

for the substitution scores. Similarly, for the gap open penalties we have

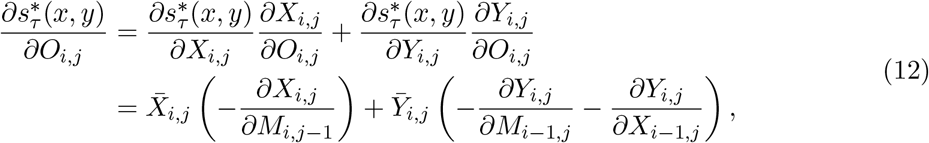

where we used that

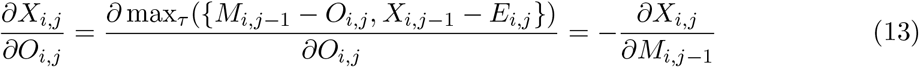

and that

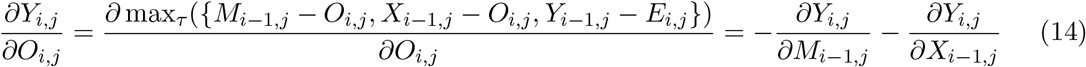

in the last step of Equation 12. Based on the same approach, the partial derivatives with respect to the gap extend penalties are given by

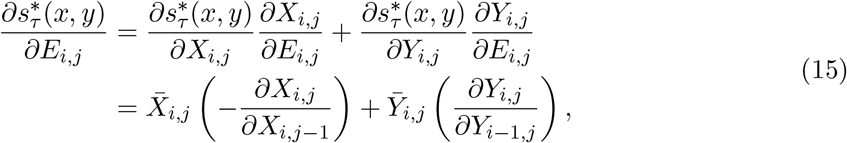

where again we used that 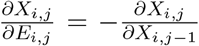 and 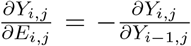, which can be shown analogously to Equations 13 and 14.

To compute the adjoint operators, we use a recursive implementation with a computation flow analogous to backtracking for the SW algorithm albeit with different update rules. Namely,

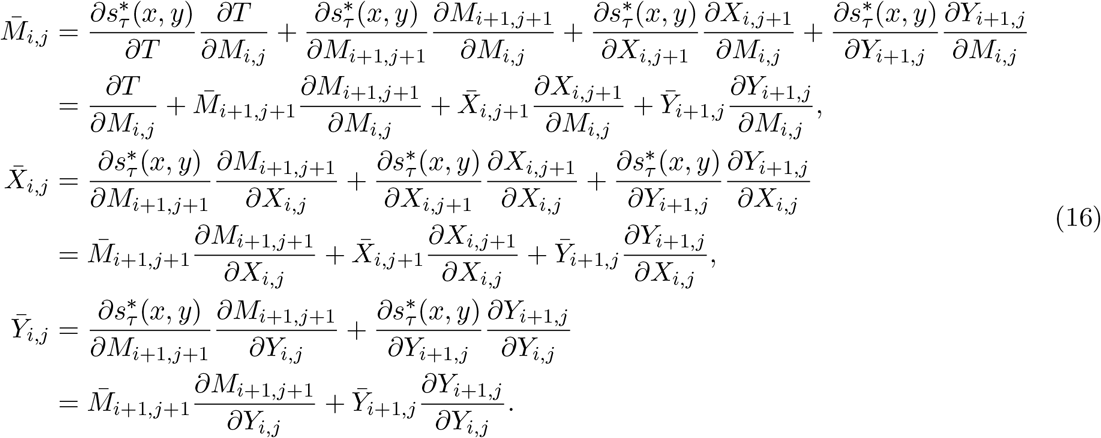

The (local) partial derivatives of the update rules in Equation 8 can all be trivially evaluated, as they are all instances of the smoothed max operator 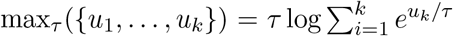 whose partial derivatives 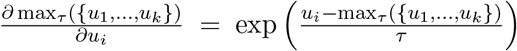 are given by the well-known *softmax* function. For instance, one may write 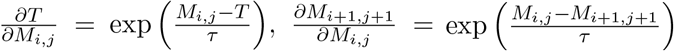 and 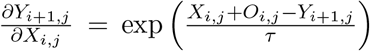. Lastly, we highlight that a careful implementation can reuse the memory used to store the values of *M_i,j_*, *X_i,j_* and *Y_i,j_* during the forward pass to also store 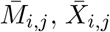 and 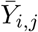 during the backward pass, roughly halving memory usage relative to a brute-force implementation.

#### S1.5 Selecting the best-performing substitution matrix baseline

We use a combination of in-distribution and out-of-distribution performance on the alignment and homology detection tasks to optimize the matrix number, gap open and gap extend penalties for each matrix family. This criterion preferentially selects versatile models that perform reliably well across all tasks under consideration over models that excel at a single task. Supposing there are *N* metrics^3^ we wish to take into account and *K* different hyperparameter settings to select from, we define an additive, self-normalized scalar summary of these *N* metrics as

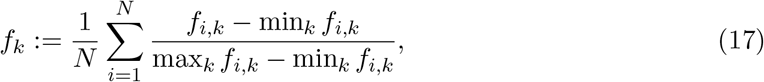

where *f_i,k_* is the value that the *k*-th hyperparameter setting attains for the *i*-th metric. The optimal hyperparameters are then simply determined as *k\** = argmax*_k_ fk*. Concretely, in this work we use *N* = 4 metrics, namely, (i) in-distribution alignment F1 score, (ii) out-of-distribution alignment F1 score, (iii) in-distribution homology detection AUPRC for remote homologs (PID < 0.1) and (iv) out-of-distribution homology detection AUPRC for remote homologs. In all cases, these metrics are computed on the validation splits, which have no overlap with the held-out test set.

### S2 Supplementary Results

#### S2.1 Ablation study

We implemented seven variations of DEDAL, each aiming to probe the effect on alignment and homology detection performance of one specific aspect of the model. These share training protocol and hyperparameters with the original DEDAL model, when applicable^4^. To further reduce the computational footprint of these experiments a single replicate was used for each ablation. Figures S5 and S6 display the alignment *F*_1_ scores and homology detection AUROC values for all approaches under study, which we discuss below.

**Figure S5:**
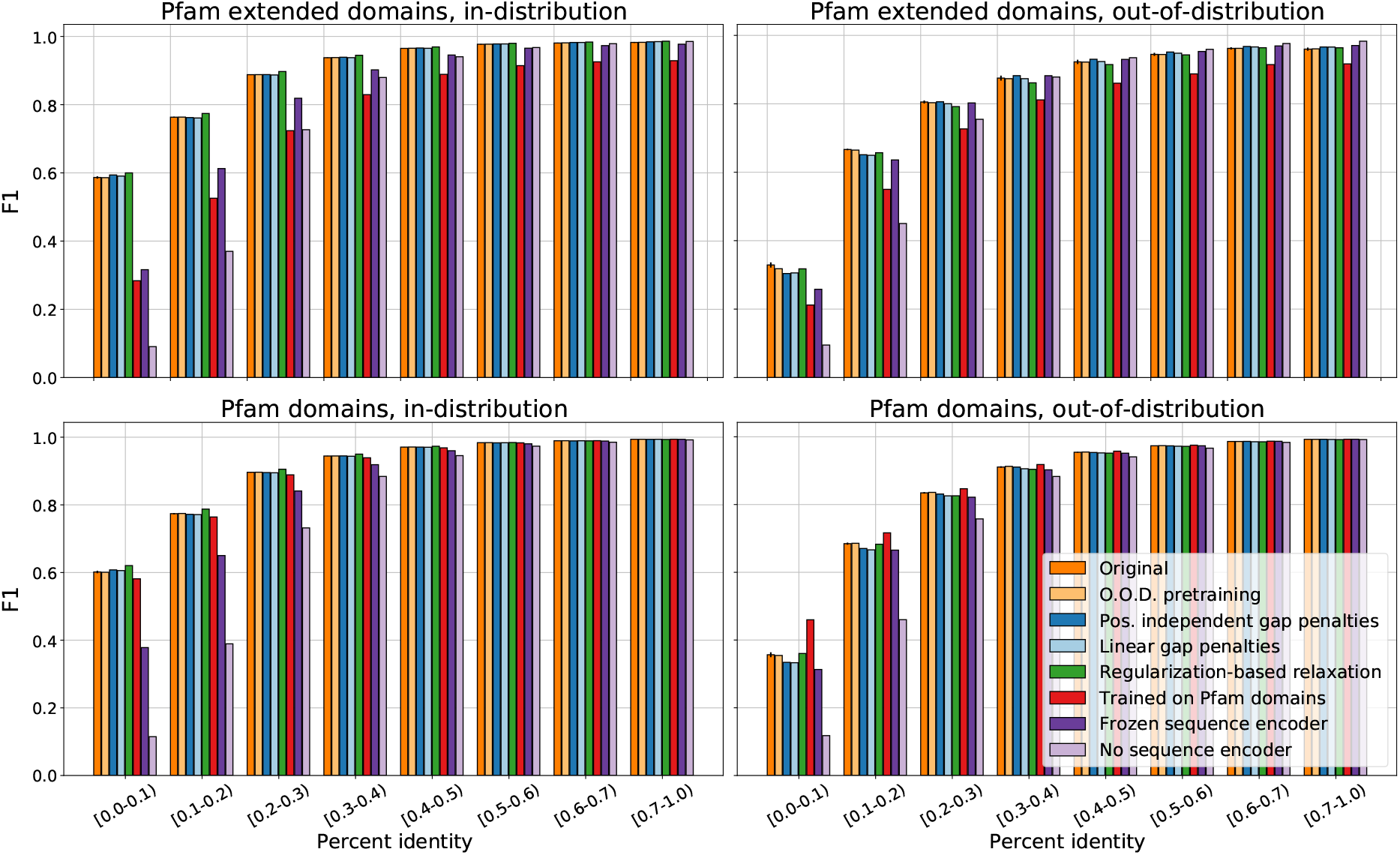
Alignment *F*_1_ score of the original DEDAL model alongside seven ablations in the in- and out-of-distribution settings (respectively, left and right columns), and for Pfam extended or raw domains (respectively, top and bottom rows). Results for the original model were averaged over 10 replicates, with error bars describing 95% confidence intervals. Ablation results are based on a single replicate for computational considerations.

##### O.O.D. pretraining

The original DEDAL model is trained on a subset of UniRef50 that contains sequences matching any Pfam families, including those in the out-of-distribution splits. Therefore, out-of-distribution performance in this regime corresponds to the case where a user wants to align sequences without any known ground-truth alignments or homologs, but which are nevertheless part of the known “protein universe”, as represented by increasingly large databases (such as UniRef50). We believe this to be the most relevant use-case in practice. However, to study the extent to which the performance of DEDAL in the out-of-distribution regime would degrade if these sequences were also excluded from pretraining, we re-trained the model using a new split of UniRef50 that simulates this more challenging scenario. To this end, we used the `hmmsearch` utility from HMMER v3.3.2 [11] to search for matches of any of the 955 Pfam clans in the out-of-distribution test set against the UniRef50 sequences at an E-value threshold of 0.01. This resulted in 1,279,217 UniRef50 sequences with significant matches to at least one Pfam family in the out-of-distribution test set, all of which were allocated to the test set for masked language modeling pretraining of this new split of UniRef50. Among the remaining 28, 882, 894 sequences with no significant matches to Pfam families in the out-of-distribution test set, we held-out an additional 90,000 and 1, 736,994 sequences for validation and test, respectively. These were chosen uniformly at random with the goal of making the new splits comparable in size to those used to train the original DEDAL model. All in all, we found this ablation to behave similarly to the original, exhibiting only a small performance degradation for remote homologs (PID < 0.1), with the alignment *F*_1_ score and homology detection AUROC dropping from 0.329 to 0.318 and from 0.910 to 0.900, respectively.

**Figure S6:**
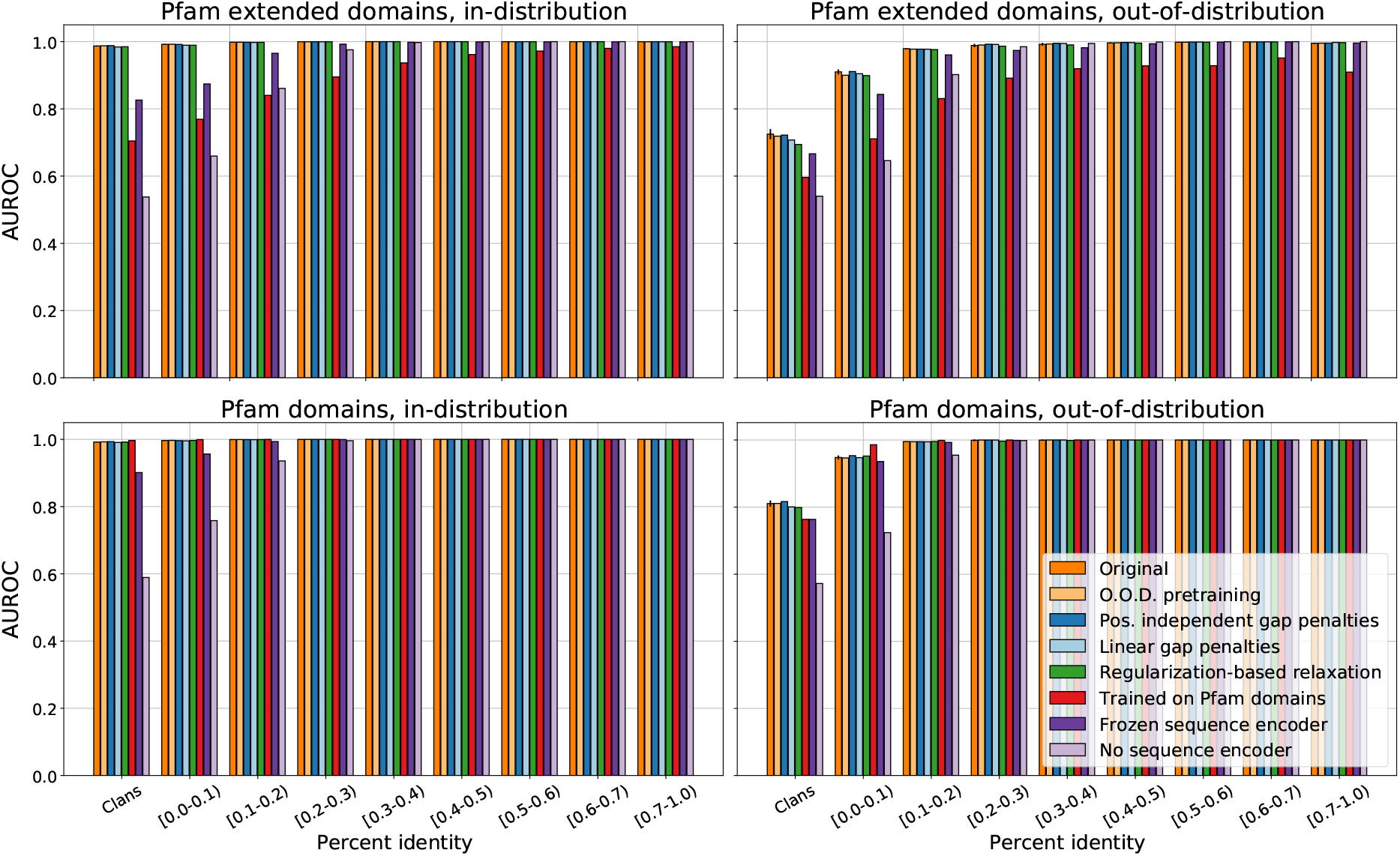
Homology detection AUROC of the original DEDAL model alongside seven ablations in the in- and out-of-distribution settings (respectively, left and right columns), and for Pfam extended or raw domains (respectively, top and bottom rows). Results for the original model were averaged over 10 replicates, with error bars describing 95% confidence intervals. Ablation results are based on a single replicate for computational considerations.

##### Position-independent gap penalties

Unlike DEDAL, traditional parameterizations of the SW algorithm use unique gap open and gap extend penalties that are shared across all positions. We tested a version of DEDAL that learns scalar, position-independent gap open and gap extend parameters instead of computing these as a function of the residue embeddings. Our findings suggest that this variant performs within the margin of error of the original model, achieving slightly inferior alignment *F*_1_ scores for remote homologs in the out-of-distribution split yet marginally superior results for sequences with PID ≥ 0.3.

##### Linear gap penalty model

DEDAL uses an affine gap penalty model, allowing it to assign different costs to opening or extending gaps in an alignment. We explored an alternative version of DEDAL based on a linear gap penalty model instead. This simplification leads to a small performance drop for remote homologs in the out-of-distribution split.

##### Regularization-based relaxation

In this work, we proposed two different techniques to relax the SW algorithm during training for end-to-end differentiability. While the final DEDAL model makes use of perturbations [7] to accomplish this, we also experimented with a variant based on smoothing via regularization [12]. Our results hint that both approaches perform comparably well, with the regularization-based version of DEDAL performing slightly better in-distribution and slightly worse out-of-distribution.

##### Trained on Pfam domains

We tried training DEDAL on the original Pfam-A seed sequences, whose ground-truth alignments are predominantly global or close to global, instead of on the extended Pfam domains we use to train DEDAL normally. As expected, this training regime enhances the performance of DEDAL when aligning Pfam domains even further, with e.g. out-of-distribution *F*_1_ scores for the smallest PID bin (< 0.1) being up to 143% superior to those of the best-performing substitution matrix baseline. However, this comes at the cost of poor generalization in scenarios where the model must predict local alignments, such as when aligning extended Pfam domains, where it only outperforms substitution matrices by a moderate margin (43% as opposed to 119% for PID below 0.1). In contrast, when DEDAL is trained on the more diverse set of extended Pfam domains, it performs consistently well both in situations requiring local alignments and those for which the sought-after alignments happen to be approximately global.

##### Frozen sequence encoder

To train DEDAL, we jointly tune the parameters of the transformer encoder network that continuously embeds input sequences and the (differentiable) alignment layer that scores and aligns them. To examine the importance of end-to-end, joint training, we first fit a transformer encoder network on a masked language modelling task and then tuned the differentiable alignment layer’s parameters while keeping the sequence encoder’s weights constant. We find this to have a mostly negative effect on in-distribution performance that is most salient for remote homologs, with e.g. alignment *F*_1_ scores dropping by up to 85% for the hardest setting (PID < 0.1).

##### No sequence encoder

Finally, we take a step further relative to the previous ablation and eliminate the sequence encoder altogether. This setting is equivalent to using our differentiable alignment layer and training scheme to learn a substitution matrix alongside scalar gap open and gap extend penalties. Unsurprisingly, we observed this to perform comparably, if not somewhat worse, than the best-performing substitution matrices from the literature.

All in all, our ablation study indicates that DEDAL is surprisingly robust to changes in many aspects of the model. Notably, the specific way in which gap penalties are parameterized appears to have only a small effect on performance, as does the choice of approach to relax the SW algorithm during training. In contrast, we found end-to-end, joint training of a flexible sequence encoder and the differentiable alignment layer to be instrumental in realizing DEDAL’s full potential.

#### S2.2 Results on the TAPE benchmark

We follow the Tasks Assessing Protein Embeddings (TAPE) benchmark [13] to probe whether using pairwise sequence alignment as supervision leads to better sequence embeddings for downstream tasks. TAPE provides standardized train, validation and test splits for five heterogeneous tasks. In a nutshell, these tasks are:

**Secondary structure:** A per-residue, 3-way (helix, strand, other) classification task, using the data from [14, 15]. It consists of 8, 678, 2,170 and 513 sequences for training, validation and testing, respectively.
**Residue-residue contacts:** This task requires predicting residue pairs that are “in contact”, defined as being less than 8Å apart. It relies on the data from [16], which provides 25,299 training sequences alongside a test set of 40 sequences (CASP12 [17]) and an additional set of 224 sequences for validation.
**Fold classification:** A per-sequence, 1,195-way classification task aiming to predict the SCOP hierarchy fold [18] each protein belongs to. Data is taken from [19] and is divided into 12,312 training, 736 validation and 718 test sequences.
**Fluorescence landscape:** A per-sequence regression task whose goal is to predict the log-fluorescence intensity of variations of a green fluorescent protein (GFP). It contains 21, 446 training, 5,362 validation and 27,217 testing sequences obtained from [20].
**Stability landscape:** Another per-sequence regression task aiming to predict a proxy for the intrinsic stability of proteins. It uses 53, 614 training, 2, 512 validation and 12, 851 testing sequences from [21].

We aim to reproduce the experimental setup used in the TAPE benchmark suite [13]. Namely, we use the same output head and losses as [13] for most tasks and fine-tune the sequence encoder while training the output head. Rather than exactly following TAPE, we chose to simplify the output heads for the secondary structure and residue-residue contact prediction tasks for computational considerations. Concretely, for secondary structure prediction, we did not employ NetSurfP-2.0 [14]. Instead, we used a simpler model consisting of a stack of two 1D convolutional layers followed by an affine mapping with output dimension three, corresponding to the number of classes (helix, strand, other). The convolutional layers have 256 and 128 filters respectively, with kernel size 10, ReLU activations and dropout with rate 0.1. Similarly, for contact prediction we simplify the RaptorX-based output head [22] used in TAPE by (i) applying linear dimensionality reduction layer that halves the dimensionality of the residue embeddings and (ii) reducing the number of ResNet blocks from 30 to 6.

For each task in the TAPE benchmark suite, we consider three different initializations for the sequence encoder: (i) random, (ii) pretrained on masked language modelling only and (iii) pretrained with DEDAL. In all cases, we use the Adam optimizer with a fixed learning rate, treated as a hyperparameter to choose from {10^−3^,10^−4^,10^−5^}, and apply early stopping to select the number of training steps. Both are tuned independently for each task to maximize the task’s corresponding metric on its validation set.

The results for all five problems are summarized in Table S2. We find that both the masked language modelling and DEDAL succeed in learning sequence representations that are transferable to downstream tasks. Indeed, initializing the model from scratch is only the superior strategy in one of the five problems, stability landscape prediction, which happens to be that which has the largest amount of training data. When it comes to comparing masked language modelling to DEDAL, our results suggest that both strategies perform similarly, with the DEDAL-based initialization being marginally better at secondary structure prediction, fold classification, and marginally worse at predicting contacts and the stability landscape.

**Table S2:** Performance on downstream tasks from the TAPE benchmark suite for different initializations of the sequence encoder: (i) random (from scratch), (ii) pretrained on masked language modelling only and (iii) pretrained on alignment, homology detection and masked language modelling (DEDAL). The secondary structure (SS) and fold classification (Folds) tasks are evaluated in terms of classification accuracy (higher is better). The protein engineering tasks (fluorescence and stability) are quantified according to the mean squared error (MSE, lower is better). Finally, the residue-residue contact prediction task (Contact) uses AUPRC for medium range contacts (higher is better). All results are averaged over 10 replicates, with the standard deviation indicated in parenthesis.

| Task | Measure | Random | Language modelling | DEDAL |
| --- | --- | --- | --- | --- |
| SS | Accuracy | 0.700 (0.001) | 0.755 (0.001) | **0.757** (0.001) |
| Contact | AUPRC | 0.158 (0.008) | **0.380** (0.020) | 0.378 (0.009) |
| Folds | Accuracy | 0.096 (0.009) | 0.232 (0.011) | **0.263** (0.011) |
| Fluorescence | MSE | 0.413 (0.132) | 0.153 (0.009) | **0.150** (0.018) |
| Stability | MSE | **0.213** (0.027) | 0.296 (0.052) | 0.309 (0.058) |

### S3 Supplementary Figures

**Figure S7:**
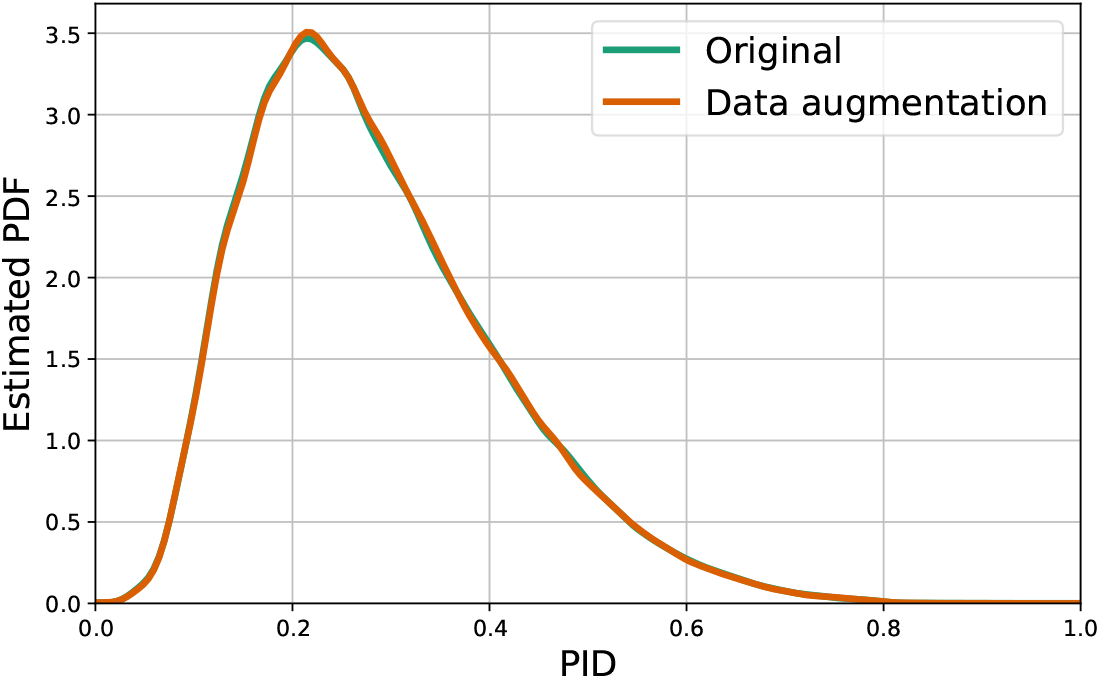
A kernel density estimate of the PID distribution for sequence pairs with and without data augmentation. 128,000 sequence pairs from the training set were used to estimate each distribution.

**Figure S8:**
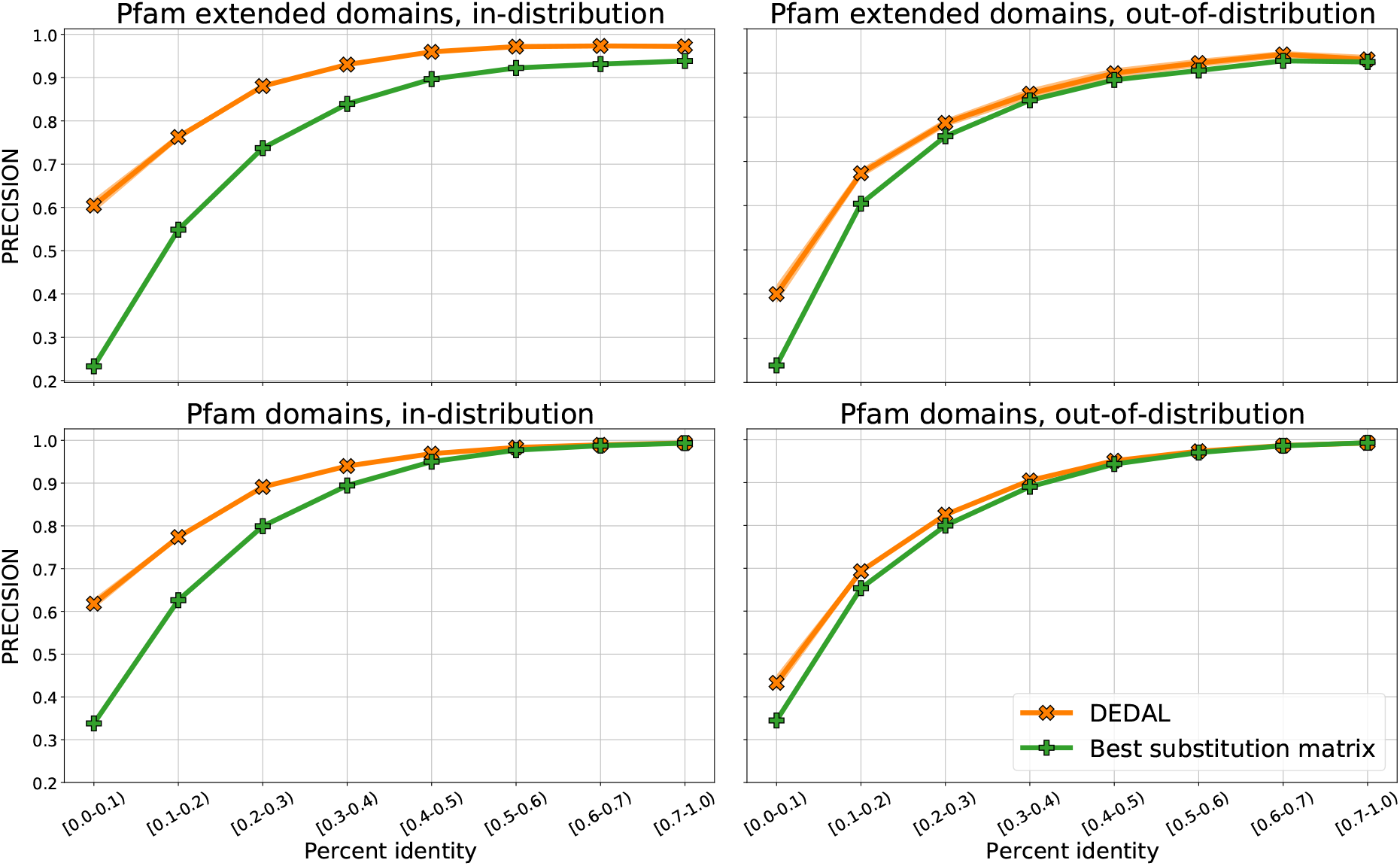
Alignment precision of DEDAL and the best-performing substitution matrix baseline, in the in- and out-of-distribution settings (respectively, left and right columns), and for Pfam extended or raw domains (respectively, top and bottom rows).

**Figure S9:**
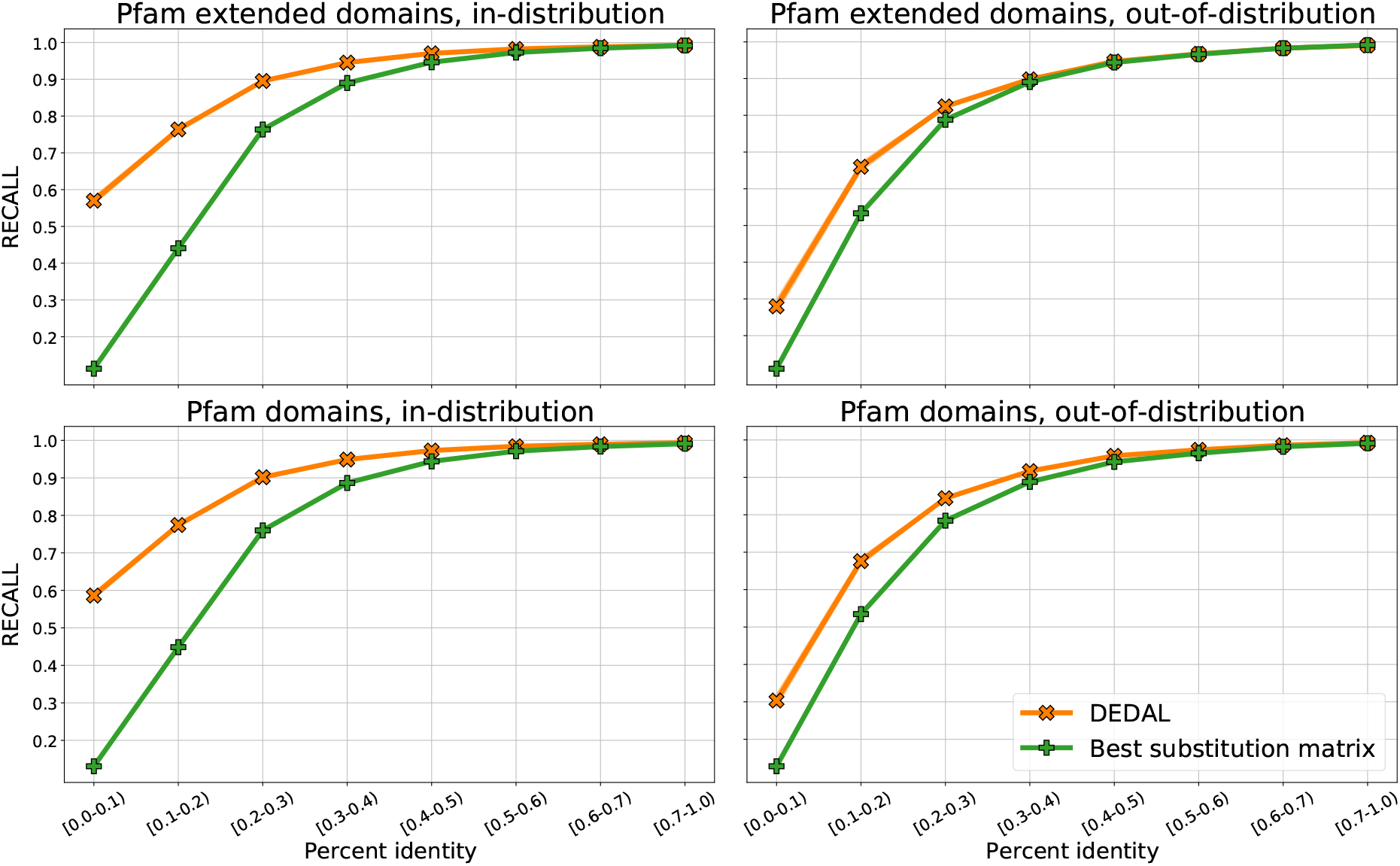
Alignment recall of DEDAL and the best-performing substitution matrix baseline, in the in- and out-of-distribution settings (respectively, left and right columns), and for Pfam extended or raw domains (respectively, top and bottom rows).

**Figure S10:**
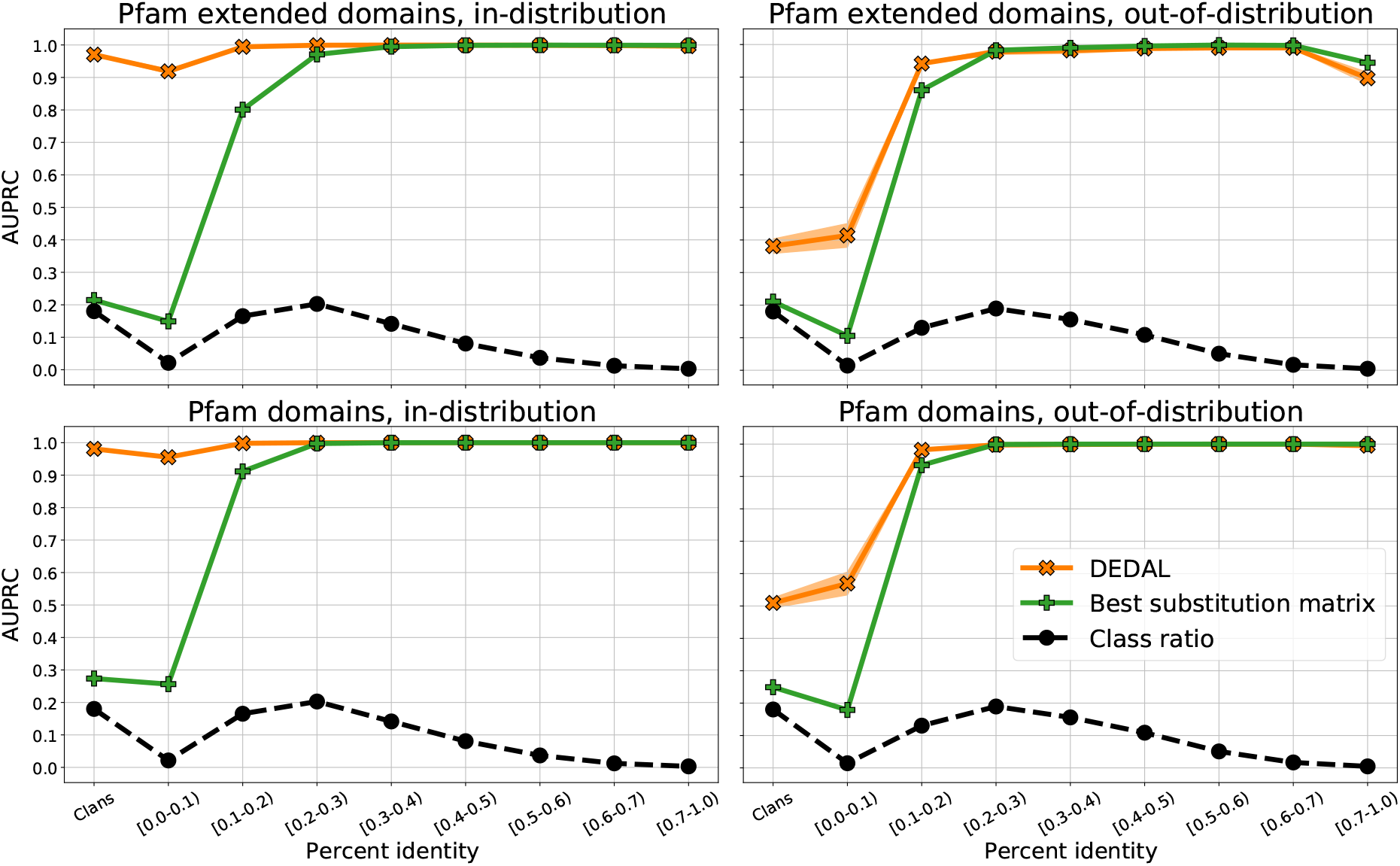
Homology detection AUPRC of DEDAL and the best-performing substitution matrix baseline, in the in- and out-of-distribution settings (respectively, left and right columns), and for Pfam extended or raw domains (respectively, top and bottom rows).

## Footnotes

1 These represent less than 0.1% of non-homologous sequence pairs.

1 As a corollary, UniProtKB sequences containing two or more Pfam-A seed domain sequences that form part of different splits may *not* be used to generate flanks.

2 In practice, we rely on the entries of the `pfamA_reg_full_significant`, `uniprot_reg_full` and `pfamA_reg_seed` tables in Pfam’s SQL database as the source of clan annotations for UniProtKB sequences.

3 For the sake of simplicity, we tacitly assume all metrics have been defined so that larger values imply better performance. In practice, we multiply the metric by minus one before normalization and aggregation whenever this is not the case.

4 Unlike for all other ablations and the original DEDAL model, we found that disabling the homology loss for the first 300,000 steps was detrimental for the performance of the ablation trained on the original Pfam domains. Thus, we omit this solely for this ablation.

## Notes

### Competing Interest Statement

All authors are employees of Google.

### Summary of Updates

Additional results and corrections in the way the benchmark for homology detection and alignment prediction is built.

